# Mechanosensitive Endothelial METTL7A Regulates Internal m^7^G mRNA Methylation and Protects Against Atherosclerosis

**DOI:** 10.1101/2025.05.22.655328

**Authors:** Tzu-Pin Shentu, Tong Wu, Zhengjie Zhou, Chih-Fan Yeh, Jiayu Zhu, Jin Li, Bernadette A. Miao, Tzu-Han Lee, Lisheng Zhang, Ru-Ting Huang, Devin Harrison, Chani J. Hodonsky, Gaelle Auguste, Aliya Husain, Matthew V. Tirrell, Clint L. Miller, Bryan Dickinson, Kai-Chien Yang, Yun Fang

## Abstract

**BACKGROUND:** Internal *N*^7^-methylguanosine (m^7^G) is a recently identified chemical modification of mammalian mRNA and a component of the epitranscriptome. While the epitranscriptome plays a key role in regulating RNA metabolism and cellular function, the specific contribution of internal m^7^G to cardiovascular health and disease remains unknown. Atherosclerosis preferentially develops at sites where disturbed blood flow activates endothelial cells, but whether internal m^7^G and its regulatory machinery influence endothelial mechanotransduction and atherogenesis is unclear.

**METHODS:** We integrated epitranscriptomic profiling, human tissue analysis, genetically modified mouse models, and targeted nanomedicine approaches to investigate the role of Methyltransferase-like protein 7A (METTL7A), a putative internal m^7^G methyltransferase, in regulating the flow-sensitive endothelial transcriptome and atherosclerosis. Vascular endothelial cells were subjected to well-defined athero-protective and athero-prone flow waveforms *in vitro* and *in vivo*. METTL7A function was assessed using RNA sequencing (RNA-seq), liquid chromatography-tandem mass spectrometry (LC-MS/MS), crosslinking immunoprecipitation sequencing (CLIP-seq), RNA stability assays, and a CRISPR-Cas-inspired RNA targeting system (CIRTS). METTL7A expression in human coronary arteries with and without atherosclerosis was evaluated by RNA-seq and immunostaining. *In vivo* atherosclerosis studies were conducted in both global and endothelial-specific *Mettl7a1* knockout mice. Endothelial METTL7A expression was restored using cationic polymer-based nanoparticles delivering CDH5 promoter-driven METTL7A plasmids or VCAM1-targeted lipid nanoparticles delivering N1-methylpseudouridine (m¹Ψ)-modified METTL7A mRNA.

**RESULTS:** Athero-protective unidirectional flow significantly induced METTL7A expression, which promoted internal m^7^G methylation of endothelial transcripts, while other major epitranscriptomic marks and cap-associated m^7^G were not affected by METTL7A. METTL7A preferentially binds to AG-enriched motifs in protein-coding mRNAs and plays a key role in regulating KLF4 and NFKBIA transcripts, enhancing their internal m^7^G and stability and supporting vascular homeostasis. In contrast, endothelial METTL7A expression was significantly reduced by disturbed blood flow and in human atherosclerotic lesions. Global or endothelial-specific loss of METTL7A exacerbated disturbed flow-induced atherosclerosis in mice, independent of serum lipid levels. Restoration of endothelial METTL7A, via nanoparticle-mediated plasmid or m^1^Ψ mRNA delivery, markedly reduced lesion formation in *Mettl7a1^⁻/⁻^* and *ApoE^⁻/⁻^* mice.

**CONCLUSIONS:** These findings establish METTL7A as a previously unrecognized mechanosensitive methyltransferase that maintains endothelial homeostasis by stabilizing key anti-inflammatory transcripts, KLF4 and NFKBIA, through internal m^7^G methylation. Loss of METTL7A disrupts endothelial function and accelerates atherogenesis in response to disturbed flow. Therapeutic restoration of endothelial METTL7A, via targeted nanoparticle-mediated gene or m^1^Ψ mRNA delivery, significantly lessens atherosclerosis. Collectively, these results uncover a novel epitranscriptomic mechanism governing vascular health and position METTL7A as a promising target for precision nanomedicine in atherosclerotic cardiovascular disease.

## INTRODUCTION

Atherosclerotic vascular complications, including myocardial infarction and ischemic stroke, remain leading causes of morbidity and mortality worldwide^1^. Atherosclerosis develops preferentially at arterial curvatures, branches, and bifurcations, where disturbed blood flow triggers endothelial inflammation and dysfunction—critical drivers of plaque initiation and progression^2–6^. In contrast, unidirectional blood flow in straight vessels promotes endothelial quiescence, providing protection against atherogenesis. Elucidating endothelial mechano-sensitive pathways activated by disturbed flow, which drive atherogenesis, and engineering precision therapeutics to modulate these pathways in inflamed endothelial cells could transform vascular wall-targeted therapies. Such approaches hold the potential to complement current pharmacological interventions, which primarily address systemic risk factors, by directly targeting disease-driving mechanisms within the vascular endothelium.

Internal chemical modifications of mRNA, collectively known as epitranscriptomes, have emerged as crucial post-transcriptional regulators of gene expression^7,8^. While most epitranscriptomic studies have focused on *N*^6^-methyladenosine (m^6^A)^9,10^, the roles of other internal mRNA modifications in human disease remain poorly understood. Among these, *N^7^-*methylguanosine (m^7^G) is a positively charged modification essential for eukaryotic mRNA 5′ capping^11^ and also occurs internally in tRNA^12^ and rRNA^13^. The METTL1-WDR4 complex catalyzes m^7^G at tRNA position 46 (m^7^G46), with dysregulation linked to cancer and brain malformation^14,15^. Recent studies identified internal m^7^G modifications in mammalian miRNA^16^ and mRNA^17,18^. Although METTL1-WDR4 was proposed to mediate mRNA m^7^G methylation, METTL1 knockdown reduced internal m^7^G levels by only 25% in HeLa cells^17^, indicating the involvement of additional regulatory mechanisms. Compared to m^6^A, the regulatory landscape of the m^7^G methylome remains largely unexplored, and its functional significance, particularly in the cardiovascular system, is unknown. Specifically, whether internal m^7^G modifications contribute to endothelial mechanotransduction and promote atherogenesis remains to be determined.

Using an m^7^G epitranscriptome toolbox, novel transgenic mouse models, human tissues, and targeted nanomedicine approaches, we demonstrate that unidirectional blood flow induces methyltransferase-like protein 7A (METTL7A) to regulate the mechanosensitive internal m^7^G methylome of endothelial mRNA, thereby promoting vascular health. In contrast, endothelial cells exposed to disturbed blood flow exhibit reduced METTL7A expression. Mechanistically, unidirectional flow upregulates endothelial METTL7A, which binds mRNA and facilitates internal m^7^G modifications.

This mechanosensitive METTL7A–m^7^G axis contributes to the endothelial transcriptome by stabilizing *KLF4* and *NFKBIA* mRNAs, both critical for maintaining vascular homeostasis. Functionally, global or endothelial-specific deletion of METTL7A exacerbates atherosclerosis in mice, whereas targeted restoration of METTL7A in activated endothelial cells via nanomedicine-based delivery significantly mitigates atherosclerosis. Together, these findings establish METTL7A-mediated internal m^7^G methylation of mRNA as a previously unrecognized molecular regulator of endothelial mechanotransduction and position endothelial METTL7A restoration as a promising therapeutic strategy for vascular diseases.

## METHODS

All studies were conducted in accordance with protocols approved by the Institutional Review Boards. All experimental animals were assigned unique identifiers to a blind experimenter to genotypes and treatment. A block randomization method was used to assign experimental animals to groups on a rolling basis to achieve an adequate sample number for each experimental condition.

### mRNA and tRNA isolation

The total RNAs were isolated from cells with TRIzol reagent (#15596026, Thermo Fisher Scientific, Waltham, MA) and then extracted using Direct-zol RNA MiniPrep (#R2050, Zymo Research, Irvine, CA). For LC-MS/MS m^7^G analysis, mRNA was extracted by two rounds of poly-A selection using Dynabeads mRNA DIRECT kit (#61006, Thermo Fisher Scientific, Waltham, MA), as previously described^17^. For tRNA isolation, small RNAs (< 200nt) were first enriched using the mirVana miRNA Isolation Kit (#AM1560, Thermo Fisher Scientific, Waltham, MA) and separated on 15% TBE-urea polyacrylamide gel electrophoresis (#EC6885, Thermo Fisher Scientific, Waltham, MA), as previously described^19^. The gel was stained with SYBR Gold (#S11494, Thermo Fisher Scientific, Waltham, MA). The specific tRNA bands (around 70-90 bp) were sliced and recovered from the gel. Further ethanol precipitation of isolated tRNA was performed to recover the tRNA.

### LC-MS/MS

To detect internal m^7^G/G level, 50 ng poly-A selected mRNA and isolated tRNAs were digested with nuclease S1 (#EN0321, Thermo Fisher Scientific, Waltham, MA) in 20 μl reaction buffer at 37^0^C for 2 hrs. Subsequently, alkaline phosphatase (#EF0651, Thermo Fisher Scientific, Waltham, MA) was added to the mixture at 37^0^C for an additional 2 hrs. To detect 5’-cap m^7^G/G level, the same amount of mRNA (50 ng) was first incubated at 95^0^C for 5 mins and cooled on ice for 2 mins.

Phosphodiesterase I (#P3243, Sigma-Aldrich, St. Louis, MO) was then added and incubated for 2 hrs at 37^0^C. Subsequently, alkaline phosphatase was added into the mixture for additional 2 hrs. After enzymatic digestion, all samples were diluted in 35 μl water and filtered with 0.22 mm filters (#SLGVR04NL, MilliporeSigma, St. Louis, MO). Mass spectrum for m^7^G, m^6^A, m^1^A, Am, Cm, Gm, A, C or G peak was indicated as previously described^17,20^. Briefly, the nucleosides retention time was 298.1-166.1(m^7^G), 282.1-150.1 (m^6^A), 282-150 (m^1^A), 282-136(Am), 258.2-112.1 (Cm), 298.1-152.1 (Gm), 268-136 (A), 244-122(C) or 284-152 (G) determined by Agilent 6410 QQQ triple136 quadrupole LC mass spectrometer (Agilent Technologies, Santa Clara, CA). The quantification of RNA modification was performed in comparison with the standard curve, obtained from pure nucleoside standards running with the same batch of samples.

### METTL7A knockdown in HAEC

METTL7A siRNA (SI00102921, Hs_DKFZP586A0522_1) was purchased from Qiagen (Qiagen, Germantown, MD). The targeting sequence (CTGGCTTTCAGAATTAACATA) is on 3’ untranslated region (3’UTR) of METTL7A. AllStars negative control siRNA (#02781, Qiagen, Germantown, MD) was used as control siRNA in the knockdown experiments. Transfection was achieved by using lipofectamine RNAiMax (#13778150, Thermo Fisher Scientific, Waltham, MA), following the manufacturer’s protocols. The cells were harvested after 48 hours after siRNA transfection. Each of siRNA (control versus siMETTL7A) was used at 20 nM to deplete METTL7A in HAEC.

### METTL7A overexpression by adenovirus transfection

The original METTL7A construct was purchased from Addgene (plasmid #62017, Watertown, MA). The METTL7A was subcloned into pReceiver M45 (GeneCopoeia, Rockville, MA) with CMV promoter and 3x HA tag at the end. Adenovirus METTL7A was generated by Vector BioLabs (Malvern, PA) using this plasmid. HAEC were transduced with 10 MOI (multiplicity of infection) of METTL7A adenovirus using GeneJammer transfection reagent (#204132, Agilent Technologies, Santa Clara, CA). Adenovirus was removed after 4 hr transfection and replenished with fresh medium. Cellular mRNAs were extracted after two days incubation and subjected to LC-MS/MS measurement or RT-qPCR to determine gene expression.

### Generation of *Mettl7a1*^−/−^*, Mettl7a1^fl/fl^* mice, and cell type–specific *Mettl7a1* conditional knockout mice

*Mettl7a1^-/-^* and *Mettl7a1^fl/fl^* C57BL/6 mice were generated using CRISPR/Cas9 genome editing. *Mettl7a1^-/-^* C57BL/6 mice were generated employing Cas9 mRNA and two single guide RNAs (sgRNA) targeting promoter region and downstream of exon 2 of *Mettl7a1*. The promoter region was 1197 bp upstream of 5’-UTR as determined by ATAC seq dataset downloaded from ENCODE (https://www.encodeproject.org/experiments/ENCSR102NGD/). Cas9 mRNA and two sgRNAs were co-injected into C57BL/6 mouse zygotes to generate mice with a deletion of *Mettl7a1*. Briefly, the pX458 vectors expressing Cas9 and sgRNA targeting *Mettl7a1* genomic sequence were generated. The two sgRNA flanking promoter region (sgRNA1) and downstream of exon2 (sgRNA2) of *Mettl7a1* were designed using CRISPR tool website (http://tools.genome-engineering.org). The resulting sequences are sgRNA1: 5’-TAGTCCTGAGAGGAGTACTG-3’ (promoter region) and sgRNA2: 5’-AGATATCCTTGAGTACAGCC-3’ (downstream of exon 2). T7 promoter sequence (AGTACTTAATACGACTCACTATAG) was then added to the Cas9 coding region and to the sgRNAs by PCR amplification. T7-Cas9 and T7-sgRNA PCR products were used as templates for *in vitro* transcription with mMESSAGE mMACHINE T7 ULTRA Transcription Kit (#AM1345, Thermo Fisher Scientific, Waltham, MA). Both the Cas9 mRNA and the sgRNAs were purified by MEGAclear Transcription Clean-Up Kit (#AM1908, Thermo Fisher Scientific, Waltham, MA). Purified Cas9 mRNA, sgRNA1 and sgRNA2 were co-injected into one-cell mouse zygotes (C57BL/6) in M2 media (Millipore Corp. MA) using a Piezo impact-driven micromanipulator. Post-injected blastocysts were transferred into the uterus of pseudopregnant female mice at 2.5 dpc. The preparation of mouse zygotes, pronuclei microinjection of Cas9 mRNA/sgRNAs, blastocysts transfer, and initial breeding of the *Mettl7a1^-/-^* animals were performed by the Transgenic Mouse Core Laboratory in National Taiwan University. PCR primers were designed to identify founders harboring a large deletion of the intended target site, as indicated by the presence of a mutant PCR amplicon (454 base pair, 5VF2+3VR4), corresponding to the expected size of the region with exons 1 and 2 deleted (Supplemental Figure 5a). The potential off-target mutageneses of CRISPR were assayed by RFLP/sequencing analysis at off-target sites predicted by CRISPR Design Tool^21^. PCR and DNA sequencing confirmed the presence of an allele with a 10.8 kb deletion encompassing exons 1 and 2 of *Mettl7a1* in one of the founders. This founder was crossed with wild-type C57BL/6 mice to obtain *Mettl7a1*^+/-^ offspring; the F1 *Mettl7a1^+/-^* progeny was then crossed to generate homozygous *Mettl7a1^-/-^* mice. F2 *Mettl7a1^-/-^* mice were born at the expected Mendelian frequency and, at baseline, showed no detectable developmental defects or structural anomalies.

*Mettl7a1^fl/fl^* mice were generated using similar CRISPR/Cas9 genome editing approaches. In addition to Cas9 mRNA and single guide RNAs (sgRNA), two single strand donor oligodeoxynucleotides (ssODNs) were designed to insert AvrII cutting site and loxP sites in the genome sequences. Both ssODNs that carried AvrII cutting site (CCTAGG) and loxP site (ATAACTTCGTATAATGTATGCTATACGAAGTTATT) were knocked-in at promoter region and downstream of exon 2 of *Mettl7a1* respectively. Briefly, the pX458 vectors expressing Cas9 and sgRNA targeting *Mettl7a1* genomic sequence were generated. The two sgRNAs flanking *Mettl7a1* were designed using CRISPR tool website (http://tools.genome-engineering.org). The resulting sequences are sgRNA1: 5’-GGTGTCGGCAGCATGCAGAG-3’ and sgRNA2: 5’-AAATATCACTGGTCCTGAGT-3’. T7 promoter sequence was then added to the Cas9 coding region and to the sgRNAs by PCR amplification. T7-Cas9 and T7-sgRNA PCR products were used as templates for *in vitro* transcription with mMESSAGE mMACHINE T7 ULTRA Transcription Kit (#AM1345, Thermo Fisher Scientific, Waltham, MA). Both the Cas9 mRNA and the sgRNAs were purified by MEGAclear Transcription Clean-Up Kit (#AM1908, Thermo Fisher Scientific, Waltham, MA). Two ssODNs carrying the AvrII cutting site (shown in Italic) and loxP site (shown in **bold**) were 5′loxP ODN: 5’-agctcagccagtgcctaccactcctgtttcaggatatgagtttttcccctcCCTAGG**ATAACTTCGTATAATGT ATGCTATACGAAGTTATT**gcatgctgccgacacctgatatgaatatcagatgg-3’ and 3′loxP ODN: 5’-ctgcctctgcctcctgagtgctgggactaaggctgaaccaccacacccact**ATAACTTCGTATAGCATACAT TATACGAAGTTAT**CCTAGGcaggaccagtgatatttgatcagaagaatccctccc-3’. Purified Cas9 mRNA, sgRNA1, sgRNA2, together with 5′loxP ODN and 3′loxP ODN were co-injected into one-cell mouse zygotes (C57BL/6) in M2 media (Millipore Corp. MA) using a Piezo impact-driven micromanipulator. Post-injected blastocysts were transferred into the uterus of pseudopregnant female mice at 2.5 dpc. The preparation of mouse zygotes, pronuclei microinjection of Cas9 mRNA/sgRNAs, blastocysts transfer, and initial breeding of the *Mettl7a1*^fl/fl^ animals were performed by the Transgenic Mouse Core Laboratory in National Taiwan University. PCR primers were designed to identify founders harboring two loxP knock-in sequences at the intended target site, as indicated by the presence of mutant PCR amplicons (5′loxP: 365 base pair, 5VF1 + 5VR1; 3′loxP: 582 base pair, 3VF1 + 3VR1, 5′loxP and 3′loxP, 381 base pair 5VF1+3VR1) (Supplemental Figure 6a). The mutant PCR amplicons were further digested by AvrII to confirm the loxP site (5′loxP: 173+192 base pair, 3′loxP: 374+208 base pair). The potential off-target mutageneses of CRISPR were assayed by RFLP/sequencing analysis at off-target sites predicted by CRISPR Design Tool. PCR1, TA cloning, and DNA sequencing confirmed the presence of an allele with successful knock-in of two lox-P sites that flanked exons 1 and 2 of *Mettl7a1* in one of the founders. This founder was crossed with wild-type C57BL/6 mice to obtain *Mettl7a1^fl/+^*offspring; the F1 *Mettl7a1^fl/+^*progeny were then crossed to generate homozygous *Mettl7a1^fl/fl^* mice. F2 *Mettl7a1^fl/fl^* mice were born at the expected Mendelian frequency and, at baseline, showed no detectable developmental defects or structural anomalies.

To generate an inducible, endothelial-specific, conditional *Mettl7a1* mouse line, *Mettl7a1^fl/fl^* mice were bred with Cdh5-Cre/ERT2 transgenic mice (provided by R. H. Adams, London, UK; to generate *Mettl7a1^ΔECKO^* transgenic mice). For activation of the Cre-ERT2 system, tamoxifen (80 mg/kg, dissolved in corn oil) was injected intraperitoneally continuously for 5 days both in control and *Mettl7a1^ΔECKO^* mice.

### Mouse lung endothelial cell (EC) isolation

Murine lung ECs isolated from *Mettl7a1^+/+^*or *Mettl7a1^-/-^* mice, were first sorted with CD31 conjugated dynabeads (#11155D, Thermo Fisher Scientific, Waltham, MA) and then subsequently sorted with ICAM2 conjugated dynabeads (#11035, Thermo Fisher Scientific, Waltham, MA). Isolated ECs were maintained in high glucose DMEM supplemented with 25 mM HEPEs, 25 mg/ml ECGS, 100 mg/ml Heparin, 1x non-essential amino acids and 20% fetal bovine serum.

### Induction of carotid atherosclerosis via hypercholesterolemia and partial carotid artery ligation (PCAL)

Hypercholesterolemia was induced in *ApoE^⁻/⁻^* mice or in wild-type mice injected via the tail vein with AAV9-PCSK9 (AAV9-HCRApoE/hAAT-D377Y-mPCSK9; 1 × 10^11^ VG; Vector Biolabs, Devault, PA), which expresses a gain-of-function mutant of PCSK9 under a liver-specific promoter. All mice were fed a high-fat diet (HFD; TD.88137, Harlan-Envigo, Indianapolis, IN) beginning either three days prior to (*ApoE^⁻/⁻^*) or on the day of (AAV9-PCSK9) injection. Carotid atherosclerosis was induced by partial ligation of the left common carotid artery (LCA) to create disturbed flow^22,23^. Mice were anesthetized with ketamine (100 mg/kg) and xylazine (10 mg/kg). A midline cervical incision was made to expose the carotid arteries, and the left external carotid artery (ECA), internal carotid artery (ICA), and occipital artery (OA) were ligated using 6-0 silk sutures, while the superior thyroid artery (STA) was left intact. Incisions were closed and mice were monitored until recovery*. ApoE^⁻/⁻^* mice remained on HFD for the duration of the study. For AAV9-PCSK9-injected mice, PCAL was performed one week post-injection. HFD continued for one additional week in *Mettl7a1^⁺/⁺^* and *Mettl7a1^⁻/⁻^* mice, and two additional weeks in *Cdh5-Cre/ERT2* and *Mettl7a1^ΔECKO^* mice. At experimental endpoints (2–3 weeks post-PCAL), mice were euthanized, and tissues were harvested following saline perfusion via the left ventricle after transection of the inferior vena cava. Carotid arteries and hearts were collected for histology or molecular analysis. Arteries were embedded in Tissue-Tek O.C.T. compound (VWR #25608-930) and cryosectioned (Leica CM3050). Carotid intima and media/adventitia RNA were isolated for qRT-PCR.

### Lipid profile analysis

Blood samples were collected by cardiac puncture under terminal anesthesia and transferred into microcentrifuge tubes containing 10 µL of EDTA (#AM9261, Thermo Fisher Scientific, Waltham, MA) to prevent coagulation. Samples were centrifuged at 4°C at 3,000 rpm for 20 minutes to separate plasma. The resulting plasma was diluted 1:10 in assay buffer, and total cholesterol levels were measured using the Amplex Red Cholesterol Assay Kit (#A12216, Thermo Fisher Scientific, Waltham, MA), following the manufacturer’s instructions.

### Oil Red O Staining for Atherosclerotic Lesion Analysis

Atherosclerosis in the ligated left carotid artery (LCA) was assessed through gross morphological evaluation and histological staining. Carotid arteries were embedded in O.C.T. compound, sectioned at 8 μm thickness using a cryostat, and serially stained with Oil Red O (#O1391, Sigma-Aldrich, St. Louis, MO) to visualize lipid-rich lesions. Images were captured using an EVOS XL Core microscope. Plaque areas were quantified using ImageJ software (NIH) by selecting Oil Red O-positive regions.

### METTL7A expression in human coronary artery tissue

For RNA-seq analysis, ischemic and non-ischemic coronary artery tissues were obtained from explanted human hearts donated for research at Stanford University under Institutional Review Board (IRB)-approved protocols, as previously described^24,25^. Briefly, coronary arteries were dissected from transplant-rejected hearts or hearts from transplant recipients, trimmed of epicardial and perivascular adipose tissue, rinsed in cold PBS, snap-frozen in liquid nitrogen, and stored at –80 °C. Non-diseased arteries were derived from donors with preserved left ventricular function (LVEF >50%). Histological classification (lesion vs. non-lesion) was performed by an experienced cardiovascular pathologist. Samples were de-identified prior to RNA processing.

Approximately 50 mg of frozen tissue was pulverized in liquid nitrogen and homogenized in QIAzol (#79306, Qiagen, Germantown, MD) using stainless steel beads in a Bullet Blender. RNA was extracted using the miRNeasy Mini Kit (#217004, Qiagen, Germantown, MD). RNA quantity and quality were assessed using Qubit 3.0 and the Agilent TapeStation; only samples with RNA Integrity Number (RIN) >5.5 and DV200 >75 were used for library construction. Libraries were prepared using the Illumina TruSeq Stranded Total RNA Gold Kit (#20020599) and barcoded with unique dual indexes (#20022371). After quality control, libraries were sequenced on an Illumina NovaSeq S4 platform (paired-end 150 bp) to a median depth of ∼100 million reads per sample. Raw reads were demultiplexed with bcl2fastq, adapter-trimmed with Trim Galore, and aligned to the human reference genome (hg38) using STAR (v2.7.3a) in two-pass mode. PCR duplicates were marked using Picard, and reads prone to mapping bias were filtered with WASP. Read counts and transcript abundances were calculated using RNA-SeQC and RSEM, with GENCODE v30 annotations. Differential gene expression analysis was performed using DESeq2, comparing either ischemic vs. non-ischemic conditions or lesion vs. non-lesion histological classifications. Age, sex, and RNA RIN were included as covariates. Genes with a Benjamini-Hochberg adjusted p-value (FDR) <0.05 and fold change ≥±50% were considered significantly differentially expressed.

To assess endothelial METTL7A protein expression in human arteries, human coronary artery tissues were obtained from deidentified patients under IRB protocol #24-0600-AM001, approved by the University of Chicago Institutional Review Board. Paraffin-embedded arterial segments were sectioned at 8 μm thickness. Hematoxylin and eosin (H&E) staining and CD31 immunohistochemistry were performed by the Human Tissue Resource Center (HTRC) at the University of Chicago. A board-certified pathologist classified arterial regions as non-lesion, atherosclerotic lesion, or atherosclerotic lesion with calcification. For METTL7A immunofluorescence, sections were deparaffinized, subjected to antigen retrieval, permeabilized, and blocked with 0.1% Triton X-100 and 5% BSA in PBS for 1 hour. Slides were incubated overnight at 4 °C with anti-METTL7A rabbit polyclonal antibody (#17092-1-AP, Proteintech, Rosemont, IL), followed by DyLight 594-conjugated secondary antibody (1:200, #35561, Thermo Fisher Scientific, Waltham, MA) for 2 hours at room temperature. Nuclei were counterstained with DAPI mounting medium (#ab104139, Abcam, Waltham, MA). Imaging was performed on an Olympus VS200 slide scanner at 40× magnification. METTL7A fluorescence intensity in CD31⁺ endothelial region was quantified by mean gray value analysis using ImageJ (NIH).

### Cell culture and application of hemodynamic flow in vitro

Human aortic endothelial cells (HAEC; CC-2535, Lonza) were cultured as previously described^26,27^. Briefly, cells were seeded in six-well plates at 100% confluence and maintained in EGM-2 medium supplemented with 4% dextran (#31392, Sigma-Aldrich, MO, USA). Cells were subjected to either unidirectional flow (UF; simulating hemodynamics found in the human distal carotid artery) or disturbed flow (DF; simulating hemodynamics in the carotid sinus) using a cone-and-plate viscometer, as previously described^26,27^. The flow device consisted of a computerized stepper motor (UMD-17, Arcus Technology, Livermore, CA) connected to a 1° tapered stainless steel cone to generate physiologically relevant shear stress patterns.

### Engineering and delivery of a CRISPR-Cas-inspired RNA targeting system

The CRISPR-Cas-inspired RNA targeting system (CIRTS) was engineered to deliver effector proteins, including epitranscriptomic regulators, to specific target transcripts^28^. Briefly, the CIRTS platform consists of a single-stranded RNA (ssRNA) binding protein, an RNA hairpin-binding protein, an effector protein, and a guide RNA (gRNA). For this study, we cloned a CIRTS plasmid encoding the human hairpin-binding protein U1A (TBP domain), a viral ssRNA-binding protein (ORF5), and human METTL7A as the effector. Guide RNAs were designed based on METTL7A binding sites in KLF4 and NFKBIA transcripts, identified by METTL7A CLIP-seq and m7G-RIP-qPCR analyses: **KLF4 mRNA-targeting guide RNA:** 5’-cccgaggggctcacgtcgttgatgtccgccaggttgaagg-3’ **NFKBIA mRNA-targeting guide RNA**: 5’-caggagatccgcctcgagccgcaggaggtgccgcgcggct-3’ CIRTS plasmids and corresponding guide RNA plasmids were delivered into human aortic endothelial cells (HAEC) using a fluorinated polyethyleneimine (F-PEI)-based nano-delivery system developed in-house. Heptafluorobutyric anhydride was reacted with PEI 1.8k (1.0 mL, 100 mg/mL in methanol) at a molar ratio of 7:1, with triethylamine as an acid scavenger. The mixture was stirred at room temperature for 48 hours, then dialyzed (MWCO = 1000 Da) against distilled water and lyophilized. Final products were characterized by 1H-NMR and 19F-NMR spectroscopy. The DNA:F-PEI molar ratio for transfection was set at 2:1. Total RNA was harvested from HAEC 48 hours post-transfection for analysis.

### Quantitative Real-Time PCR (qPCR)

For human aortic endothelial cells (HAEC), total RNA was extracted and reverse-transcribed using the High-Capacity cDNA Reverse Transcription Kit (#4368814, Thermo Fisher Scientific, Waltham, MA). For mouse samples, intimal and medial/adventitial RNA from individual carotid arteries was isolated and reverse-transcribed using Maxima H Minus Reverse Transcriptase (#EP0751, Thermo Fisher Scientific, Waltham, MA). Transcript concentrations were determined using external standard curves on a LightCycler 480 II system (Roche). Human and mouse qPCR primers are listed in Table S1. Absolute transcript levels were normalized to the geometric mean of β-actin, GAPDH, and ubiquitin expression.

### METTL7A CLIP-seq

METTL7A CLIP-seq experiments were conducted based on a previously reported protocol^17^ with minor modifications. HA-tagged METTL7A was overexpressed in HAEC via adenoviral infection, and cells were crosslinked using 0.15 J/cm² of 365 nm UV light twice on ice to minimize heating. Cells were collected with lifters, lysed with lysis buffer (4× the pellet volume), and the supernatant was clarified. The lysate was incubated with RNase T1 (#EN0541, Thermo Fisher Scientific, Waltham, MA) at 22°C for 15 minutes and quenched on ice. The supernatant was then incubated with anti-HA antibody-conjugated Protein A Dynabeads (6 μg antibody to 30 μL beads; #10001D, Thermo Fisher Scientific, Waltham, MA) overnight at 4°C under rotation to immunoprecipitate METTL7A–RNA complexes. Beads were washed three times with 1 mL immunoprecipitation wash buffer (50 mM Tris-HCl, 300 mM KCl, 0.05% NP-40, 0.5 mM DTT, supplemented with protease inhibitor cocktail) and then transferred to new tubes for five additional washes with high-salt buffer (50 mM Tris-HCl, 500 mM KCl, 0.05% NP-40, 0.5 mM DTT). The bound complexes were digested with proteinase K, and RNAs were purified using the RNA Clean & Concentrator-5 kit (#R1013, Zymo Research, Irvine, CA). Eluted RNA was treated with T4 polynucleotide kinase (#EK0031, Thermo Fisher Scientific, Waltham, MA) in 10× T4 PNK buffer at 37°C for 30 minutes, followed by re-purification. Small RNA libraries were prepared using the NEBNext Small RNA Library Prep Set for Illumina (#E7560, New England BioLabs, Ipswich, MA) and sequenced on an Illumina NextSeq 500 platform with single-end 80 bp reads.

### Analysis of CLIP-seq Data and Motif Identification

Raw sequencing reads were processed using Trim Galore (https://www.bioinformatics.babraham.ac.uk/projects/trim_galore/) to remove adapter sequences and low-quality reads (<20 nucleotides). Cleaned reads were aligned to the human reference genome (hg38) using HISAT2 (version 2.1.0), and unmapped reads were removed with SAMtools (version 1.6.0). Enriched peaks in immunoprecipitated (IP) samples compared to input controls were identified using OmniCLIP. Peaks with p-values ≤ 0.05 from each biological replicate were annotated with PAVIS (https://manticore.niehs.nih.gov/pavis2/), and results were merged based on transcript type and structural elements. To increase detection sensitivity, peak calling was also performed using Pyicoclip from the Pyicoteo Tools Suite (https://bitbucket.org/regulatorygenomicsupf/pyicoteo/src/pyicoteo/pyicoteolib/), focusing on exon regions (-region-magic option). Pyicoclip was applied to reads between 40–80 nt, and peak significance was determined using a modified false discovery rate (FDR) method. Enriched peaks were further filtered with Pyicoenrich, allowing pseudocounts to retain regions detected only in IP samples. Genes of interest were selected based on Pyicoenrich output. Motif analysis was performed using HOMER (version 4.10.0; http://homer.ucsd.edu/homer/motif/) with the RNA-specific (-rna) option. IP-derived sequences were used as target input, and input-derived sequences served as the background. The most significantly enriched motif was based on p-value.

### RNA-seq and sequence data analysis in HAEC

RNAs isolated from HAEC treated with siMETTL7A or siCtrl and under 24-hr unidirectional flow were digested with DNAase in column using Direct-zol RNA MiniPrep (#R2050, Zymo Research, Irvine, CA). The quality of RNAs was assessed by 2100 Bioanalyzer Instrument (Agilent Technologies, Santa Clara, CA) and sent for sequencing on Illumina HiSeq 4000 with pair-end 100 bp read length. Raw Read text file was stored in the FASTQ format. Quality of reads was assessed using fastQC. Reads were aligned to the reference genome (human hg38) using hisat2 (version 2.1.0). Transcriptome were then assembled using Cufflinks (version 2.2.1), merged by Cuffmerge (version 2.2.1), and differentially expressed genes were identified using Cuffquant (version 2.2.1) followed by Cuffdiff (version 2.2.1). FDR 0.1 was used as cut-off for differentially expressed genes. The sequencing data have been deposited in the Gene Expression Omnibus (GEO) under accession number GSE235521. The data can be accessed using the following token: mdklsiumxvsbhyn.

### Gene Ontology Analysis

Gene ontology (GO) analysis of differentially expressed genes from the RNA-seq dataset was performed using Ingenuity Pathway Analysis (IPA; https://digitalinsights.qiagen.com)^29^ and the Database for Annotation, Visualization and Integrated Discovery (DAVID; https://david.ncifcrf.gov)^30^. GO terms were sorted by the category GOTERM_BP_FAT. Terms with *p* < 0.05 were considered statistically significant.

### Internal m⁷G RNA immunoprecipitation and quantitative PCR (m⁷G-RIP-qPCR)

To assess internal m⁷G modification of mRNA, 2 μg of total RNA was subjected to two rounds of poly(A) selection followed by enzymatic decapping using acid pyrophosphatase (# C-CC15011H, CellScript, Madison, WI). The decapped mRNA (500–600 ng) was purified using the RNA Clean & Concentrator-5 kit (# R1013, Zymo Research, Irvine, CA). m⁷G-modified RNAs were immunoprecipitated using 10 μg of anti-m⁷G antibody (# RN017M, MBL, Woburn, MA) conjugated to 100 μl of protein G Dynabeads (# 10003D, Thermo Fisher Scientific, Waltham, MA). The reaction was incubated overnight at 4 °C with gentle rotation. The next day, the RNA–antibody–bead complexes were washed five times with RIP buffer (100 mM Tris-HCl, 1.5 M NaCl, 1% NP-40) and transferred to a new tube. To competitively elute m⁷G-bound RNA, the beads were incubated in 100 μl RIP buffer containing 2 μl RNaseOUT (# 10777019, Thermo Fisher Scientific, Waltham, MA) and 6.7 mM 7-methyl-GTP (# M6133, Sigma-Aldrich, St. Louis, MO) at 37 °C for 30 minutes. RNA was then purified using the RNA Clean & Concentrator-5 kit. Purified RNA was reverse-transcribed using Maxima H Minus Reverse Transcriptase (# EP0751, Thermo Fisher Scientific, Waltham, MA).

Quantitative PCR was performed to determine transcript enrichment. Absolute transcript levels of KLF4 and NFKBIA in m⁷G-IP samples were normalized to matched input RNA controls.

### In vitro m⁷G methylation assay

HA-tagged METTL7A protein was purified from HEK293T cells transfected with METTL7A expression plasmids (pReceiver M45, GeneCopoeia, Rockville, MD) using Lipofectamine 2000 (# 11668019, Thermo Fisher Scientific, Waltham, MA). After 48 hours, cells were harvested and lysed, and METTL7A protein was isolated using anti-HA magnetic beads (6 μl antibody in 50 μl bead slurry; # 10001D, Thermo Fisher Scientific, Waltham, MA). Protein elution was performed using HA peptide (# I2149, Sigma-Aldrich, St. Louis, MO), and eluted proteins were verified by SDS-PAGE. In vitro transcription of human KLF4 and NFKBIA mRNAs was carried out using the mMESSAGE mMACHINE T7 kit (# AM1344, Thermo Fisher Scientific, Waltham, MA). The methylation reaction was adapted from a previously published method^17,20^, with modifications. Briefly, 2 μg of mRNA was incubated with 2 μg of HA-tagged METTL7A in a 160 μl reaction containing 16 μl of 10 mM deuterated S-adenosyl methionine (d₃-SAM; Cat# D4093, CDN Isotopes, Canada). The reaction was performed at 16 °C overnight.

Following incubation, RNA was purified via acid phenol-chloroform extraction and RNA Clean & Concentrator kit (#R1013, Zymo Research, Irvine, CA). 150 ng of purified RNA was digested with 10 U of nuclease S1 (# EN0321, Thermo Fisher Scientific, Waltham, MA) for 2 hours at 37 °C, followed by alkaline phosphatase (#EF0652, Thermo Fisher Scientific, Waltham, MA) treatment for an additional 2 hours. Final products were diluted in 35 μl of nuclease-free water and filtered using a 0.22 μm membrane (# SLGVR04NL, MilliporeSigma, St. Louis, MO). Detection of methylated nucleosides was performed by LC-MS/MS. The retention time of d₃-m⁷G was monitored at 298.1 → 169.1 m/z. Deuterated m⁷G (# D8061, CDN Isotopes) was used as the internal standard.

### Immunofluorescence staining

Cryosections of mouse carotid arteries were fixed in 4% paraformaldehyde (PFA) in PBS, followed by permeabilization and blocking in PBS containing 1% bovine serum albumin (BSA) and 0.1% Triton X-100. Sections were incubated overnight at 4°C with primary antibodies against METTL7A (1:100), KLF4 (1:100), NFKBIA (1:100), or FLAG (1:100). After washing, sections were incubated for 1 hour at room temperature with fluorophore-conjugated secondary antibodies: Alexa Fluor 594-conjugated anti-rabbit IgG or Alexa Fluor 647-conjugated anti-mouse IgG (1:400; #A-11012 or #A-21239, Thermo Fisher Scientific, Waltham, MA). Nuclei were counterstained with DAPI. Fluorescence images were acquired using a Zeiss 3i Marianas spinning disk confocal microscope equipped with SlideBook acquisition software, using a 40×/1.3 NA oil-immersion Plan-Apochromat objective.

### Actinomycin D chase assay

To assess mRNA stability, human aortic endothelial cells (HAEC) were subjected to either siRNA-mediated knockdown or adenovirus-mediated overexpression. Transcription was halted by treating the cells with 5 μg/mL actinomycin D (#A1410, Sigma-Aldrich, St. Louis, MO). Total RNA was extracted at designated time points post-treatment using Direct-zol RNA MiniPrep (#R2072, Zymo Research, Irvine, CA), followed by reverse transcription into cDNA using the High-Capacity cDNA Reverse Transcription Kit (#4374967, Thermo Fisher Scientific, Waltham, MA). Quantitative real-time PCR (qPCR) was performed as previously described. The relative abundance of each transcript was normalized to its level at time zero to determine the decay rate.

### Western Blotting

Protein lysates were prepared from cells using cold RIPA buffer (#R0278, Sigma-Aldrich, St. Louis, MO) supplemented with protease and phosphatase inhibitors (#4906837001, Roche, Indianapolis, IN). Protein concentrations were measured using the BCA Protein Assay Kit (#23227, Thermo Fisher Scientific, Waltham, MA). Equal amounts of protein were separated by SDS-PAGE and transferred to PVDF membranes. Membranes were incubated with primary antibodies: anti-METTL7A (1:1000), anti-KLF4 (1:1000), anti-NFKBIA (1:1000), anti-HA (1:1000), anti-α-tubulin (1:1000), or anti-β-actin (1:1000), followed by HRP-conjugated anti-rabbit IgG (#31464, Thermo Fisher Scientific, Waltham, MA) or anti-mouse IgG (#401253, MilliporeSigma, St. Louis, MO) secondary antibodies (1:4000). Detection was performed using ECL substrate (#34579 or #34096, Thermo Fisher Scientific, Waltham, MA) and imaged with the ChemiDoc MP Imaging System (Bio-Rad). Band intensities were quantified using ImageJ software (NIH) and normalized to loading controls.

### In vivo endothelial mETTL7A overexpression via CDH5-METTL7A Plasmids

Plasmid delivery was performed via one or two intravenous injections of a DNA– polymer complex consisting of 50 μg of endotoxin-free plasmid DNA and a cationic transfection reagent (#R0531 TurboFect, Thermo Fisher Scientific, MA, USA), following previously established protocols ^27,31^. The plasmid encoded METTL7A under the control of the endothelial-specific VE-cadherin (CDH5) promoter^32^. A control plasmid lacking a functional gene was used as a negative control. Injections were administered via the tail vein.

### In vivo endothelial METTL7A restoration via VCAM1-targeted lipid nanoparticles delivering m1Ψ-Modified mRNA

To restore METTL7A expression in inflamed endothelial cells, we employed N1-methylpseudouridine (m1Ψ)-modified METTL7A mRNA encapsulated in lipid nanoparticles functionalized for VCAM1 targeting, as previously described^33^. The self-assembled lipid nanoparticles consisted of dioleoylphosphatidylethanolamine (DOPE), lipid-grafted polyamidoamine dendrimer (G0-C14), cholesterol, and 1,2-distearoyl-sn-glycero-3-phosphoethanolamine-poly(ethylene glycol) (DSPE-PEG), along with the mRNA payload as described^33^. To enable active targeting, a VCAM1-binding peptide (VHPKQHR) was conjugated to DSPE-PEG, facilitating selective delivery to inflamed endothelial cells. ApoE^⁻/⁻^ mice or AAV9-PCSK9-injected mice (8–10 weeks old) were fed a high-fat diet (HFD) and subjected to partial carotid artery ligation (PCAL). Mice were intravenously administered VCAM1-targeting nanoparticles encapsulating METTL7A mRNA. Nanoparticles carrying non-translatable mutant METTL7A mRNA were used as controls.

### Antibodies

The antibodies used in this study include anti-METTL7A (#TA809878, OriGene, Rockville, MD; #ab12801, Abcam, Waltham, MA; and #17092-1-AP, Proteintech, Rosemont, IL), anti-m7G (#RN017M, MBL, Woburn, MA), anti-HA (#ab9110, Abcam, Waltham, MA), anti-α-tubulin (#3873, Cell Signaling Technology, Danvers, MA), anti-β-actin (#ab6276, Abcam, Waltham, MA), anti-KLF4 (#ab106629, Abcam, Waltham, MA), anti-NFKBIA (#4814 and #4812, Cell Signaling Technology, Danvers, MA), and anti-Flag (#8146 and #14793, Cell Signaling Technology, Danvers, MA).

### Primers

The list of used primers was listed in the Supplemental Table 1.

## Results

### Unidirectional flow-induced METTL7A mediates the mechanosensitive internal m^7^G methylome of endothelial mRNA

To determine the endothelial epitranscriptome as a function of well-defined mechanical stimuli and identify mechanosensitive endothelial methyltransferases, we conducted liquid chromatography-tandem mass spectrometry (LC-MS/MS) and RNA-seq in human aortic endothelial cells (HAEC) subjected to complex hemodynamics (Figure 1a). These flow waveforms mimicked the unidirectional blood flow measured in human distal carotid artery resistant to atherosclerosis and the disturbed blood flow measured in human carotid sinus prone to atherosclerosis^34^. LC-MS/MS demonstrated that global internal m^6^A, the most abundant and well-characterized mRNA modification^35^, is not altered in endothelium by hemodynamics (Supplemental Figure 1a), but internal m^7^G of endothelial mRNA was significantly increased by unidirectional flow (UF) when compared to disturbed flow (DF) (Figure 1b). Although endothelial internal m^7^G was regulated by hemodynamics, cap-m^7^G of endothelial mRNA (Figure 1c) and m^7^G in endothelial tRNA were not significantly changed by hemodynamics (Supplemental Figure 1b). RNA-seq (absolute fold change >2; false discovery rate <5%) identified that methyltransferase-like 7A (*METTL7A*), a candidate methyltransferase, was markedly upregulated by unidirectional flow when compared to disturbed flow.

**Figure 1.**
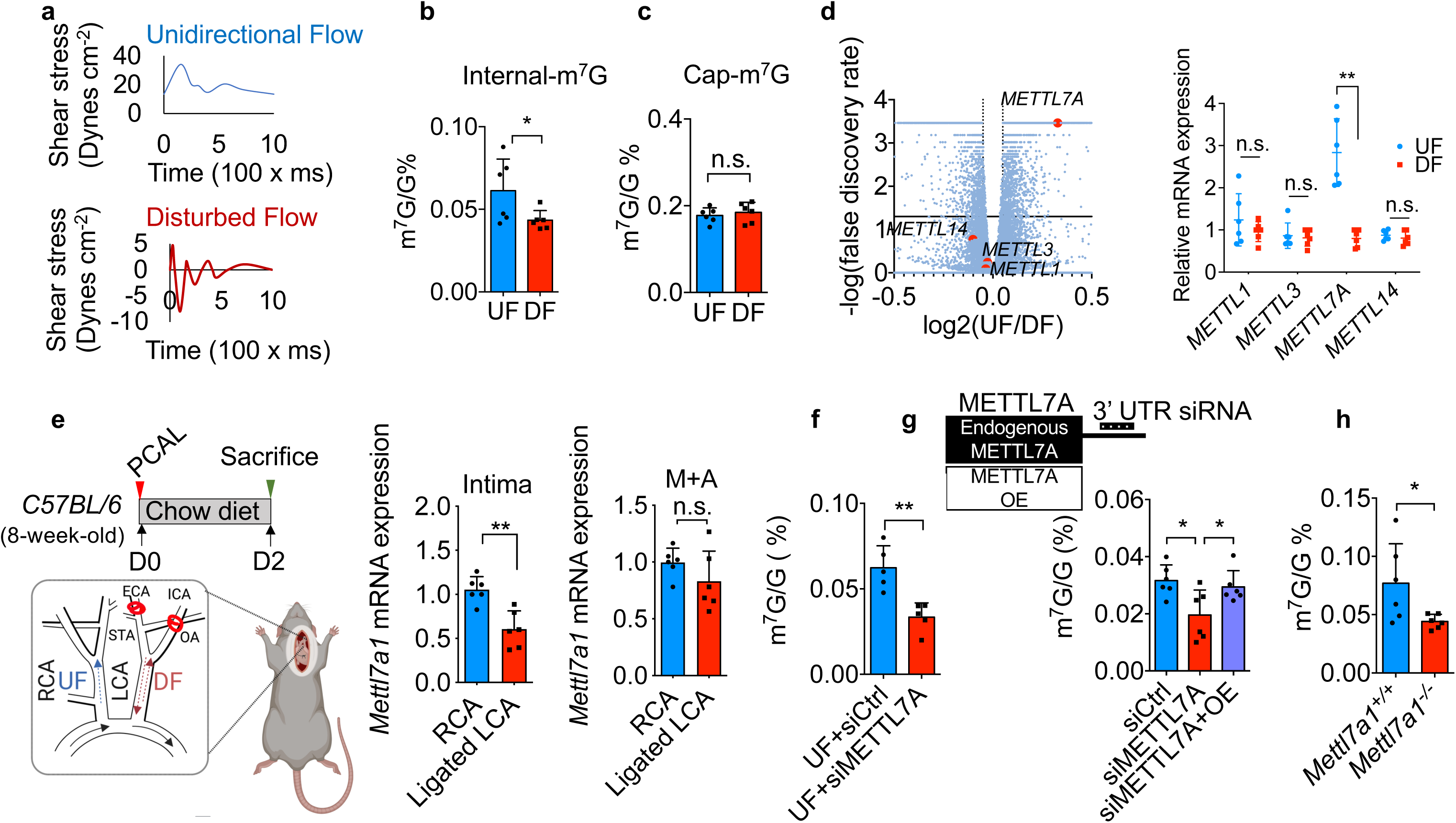
Unidirectional flow-induced METTL7A mediates internal m^7^G methylome of endothelial mRNA. **(a)** *In vitro* waveforms recreate athero-protective unidirectional flow (UF) mimicking *in vivo* hemodynamics in human distal carotid artery or athero-prone disturbed flow (DF) mimicking hemodynamics in human carotid sinus. **(b and c)** LC-MS/MS detected a significant increase of internal m^7^G but not in the cap-m^7^G of mRNA in HAEC subjected to 24-hr UF compared to cells exposed to 24-hr DF. (n = 6 biological replicates; *p* values were obtained using two-tailed Student’s t-test using GraphPad Prism. \**p*≤0.05 and n.s. indicates non-significance (*p*>0.05)). **(d)** Whole-genome RNA-seq in HAEC, illustrated by a volcano plot, detected a significant increase of endothelial *METTL7A* but not *METTL1*, *METTL3*, and *METTL14* by UF. UF-induced endothelial *METTL7A* but not *METTL1*, *METTL3*, and *METTL14* was validated by real-time PCR (n = 6 biological repeats, \*\**p*≤0.01 and n.s. indicates non-significance (*p*>0.05)). **(e)** 48-hr acute disturbed flow in the partially-ligated left carotid artery markedly reduced *Mettl7a1* transcripts in endothelium-enriched intima but not in non-endothelial media/adventitia (M+A) (n = 6 biological repeats, \*\**p*≤0.01 and n.s. indicates non-significance (*p*>0.05)). Experiments were performed on 8-week-old male *C57BL/6* mice. PCAL: partial carotid artery ligation, LCA: left carotid artery, RCA: right carotid artery, ECA: external carotid artery, ICA: internal carotid artery, OA: occipital artery and STA: superior thyroid artery. Different flow patterns were indicated in the LCA (DF) and RCA (UF). **(f)** Suppression of elevated METTL7A by siRNA in HAEC under UF significantly reduced mRNA internal m^7^G quantified by LC-MS/MS (n = 5 biological replicates, *\*\*p*≤0.01). **(g)** Reduced internal m^7^G in METTL7A-knockdowned HAEC was restored by *METTL7A* mRNA replenishment via overexpression (OE) of *in vitro* transcribed *METTL7A* mRNA without the 3’UTR targeted site by siMETTL7A (n = 6 biological repeats, \**p*≤0.05). **(h)** LC-MS/MS detected a significant decrease of mRNA internal m^7^G in lung vascular endothelial cells isolated from *Mettl7a1^-/-^* mice compared to endothelial cells isolated from wild-type *Mettl7a1^+/+^* mice (n = 6 biological repeats, \**p*≤0.05).

Meanwhile, RNA-seq and real-time qPCR data showed that hemodynamics had no significant effect on endothelial mRNA levels of *METTL1*, *METTL3*, and *METTL14,* previously identified mRNA methyltransferases (Figure 1d).

To determine the flow-sensitivity of endothelial METTL7A *in vivo*, we conducted partial carotid artery ligation (PCAL) in mice^23^. Three of the four caudal branches of the left common carotid artery (LCA) were ligated to introduce acute oscillatory disturbed flow in LCA while undisturbed and unidirectional blood flow was maintained in the right common carotid artery (RCA) (Figure 1e). A 57% mRNA reduction of endothelial *Mettl7a1* (the mouse homolog of human *METTL7A*) was detected in ligated LCA compared with RCA 48 hours after partial ligation (Figure 1e). No change of *Mettl7a1* mRNA was detected in the smooth muscle cell-enriched media/adventitia in the ligated LCA (Figure 1e). Concordant with the *in vitro* studies, acute DF *in vivo* had no effect on mRNA levels of *Mettl1*, *Mettl3*, and *Mettl14* (Supplemental Figure 1c-e) in the endothelium or media/adventitia of ligated LCA. *METTL7A* knockdown by siRNA in HAEC under UF had no effect on expression of *METTL1*, *METTL3*, and *METTL14* (Supplemental Figure 2). These results suggested the flow-sensitive nature of METTL7A in the endothelium, which contrasted with other methyltransferases.

*METTL7A* suppression with siRNA (Supplemental Figure 2a) significantly reduced internal m^7^G methylation (Figure 1f), pointing to a critical role of METTL7A in mediating the mechanosensitive endothelial internal m^7^G methylome. In contrast, the ratios of m^6^A/A, cap-m^7^G/G (Supplemental Figure 3b), Cm/C, Am/A, m^1^A/A, and Gm/G (Supplemental Figure 3c) of endothelial mRNA were not affected by *METTL7A* knockdown (Supplemental Figure 3a). Since our *METTL7A* siRNA targeted the *METTL7A* 3’UTR, we replenished endogenous *METTL7A* mRNA by introducing *in vitro* transcribed *METTL7A* mRNA (without 3’UTR, labeled as METTL7A overexpression/ OE) to *METTL7A*-depleted HAEC, which effectively restored the internal m^7^G of endothelial mRNA (Figure 1g). In addition, adenovirus-mediated overexpression of HA-tagged METTL7A protein in HAEC markedly increased internal m^7^G but not m^6^A of endothelial mRNA (Supplemental Figure 4). Taken together, these *in vivo* and *in vitro* results demonstrate that the internal m^7^G methylome of endothelial mRNA is dynamically regulated by mechanical forces and mediated by flow-sensitive expression of METTL7A.

### Mechanosensitive endothelial METTL7A lessens atherosclerosis

To determine the causal function of *Mettl7a1 in vivo*, we generated a new mouse line (*Mettl7a1^-/^*^-^) in which *Mettl7a1* is globally deleted using CRISPR/Cas9 genome editing (Supplemental Figure 5a). *Mettl7a1^-/-^* mice were born at the normal Mendelian ratio. Consistent with the *in vitro* results, we detected a marked reduction in internal m^7^G (Figure 1h) but not m^6^A (Supplemental Figure 5b) in mRNAs of lung endothelium isolated from *Mettl7a1^-/-^*mice compared to *Mettl7a1^+/+^* mice. We then induced carotid atherosclerosis, which occurs due to local DF and systemic hypercholesterolemia, with partial carotid artery ligation and expression of proprotein convertase subtilisin/kexin type 9 (PCSK9) mediated by adeno-associated virus 9 (AAV9)^23^. DF-induced atherosclerosis in the ligated LCA was significantly increased in PCSK9-AAV9-injected *Mettl7a1^−/−^* mice compared to PCSK9-AAV9-injected *Mettl7a1^+/+^* (Figure 2a) while serum cholesterol levels remained the same in these two mouse lines (Supplemental Figure 5c), demonstrating an athero-protective, lipid-independent role of *Mettl7a1 in vivo*.

**Figure 2.**
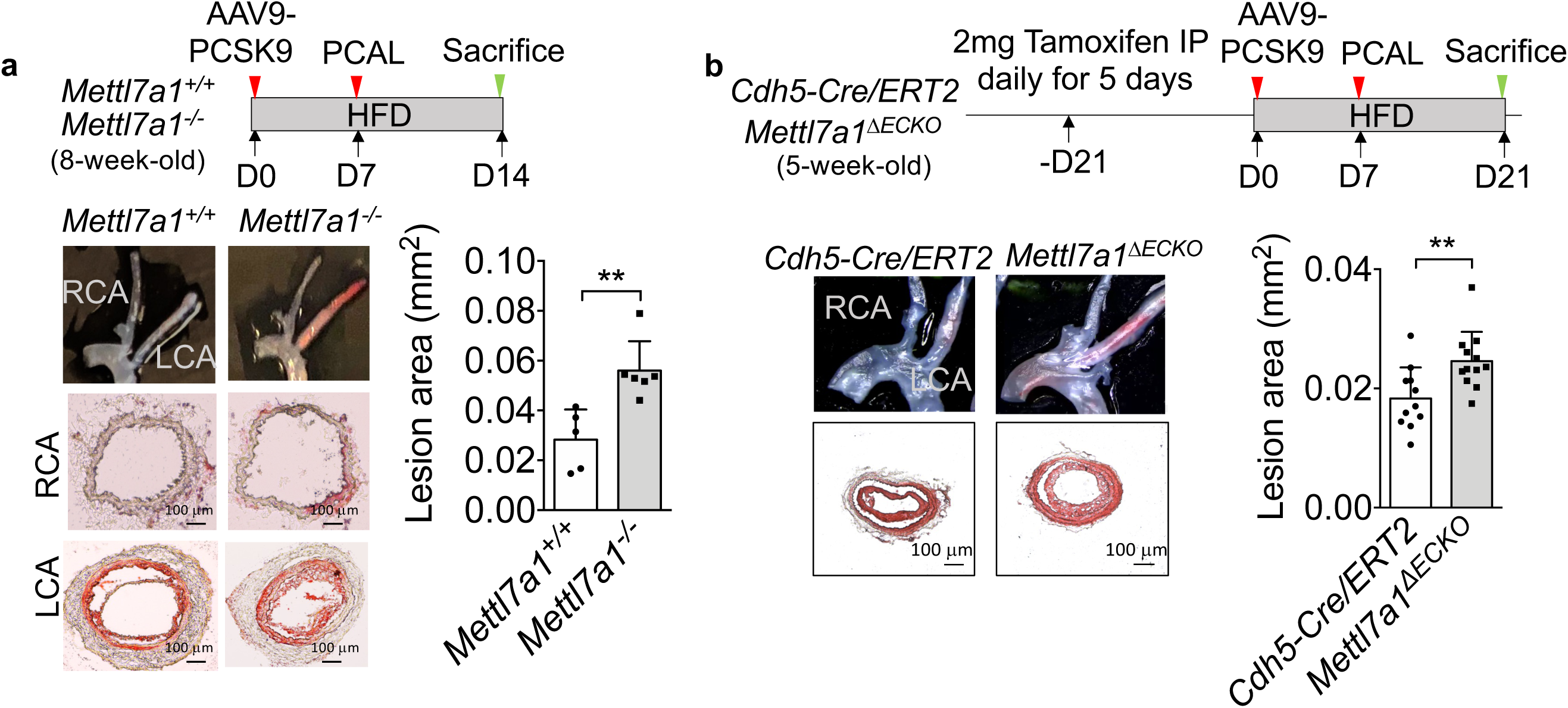
Global and endothelial-specific deletion of Mettl7a1 exacerbates atherosclerosis in mice. **(a)** Disturbed flow-induced atherosclerosis in the ligated carotid artery was significantly increased in hypercholesterolemic *Mettl7a1^-/-^* mice compared to hypercholesterolemic *Mettl7a1^+/+^* controls (n = 5 for *Mettl7a1^+/+^* mice; n = 6 for *Mettl7a1^-/-^* mice, *p* values were obtained using two-tailed Mann-Whitney U test using GraphPad Prism. *\*\*p* ≤ 0.01). **(b)** Disturbed flow-induced atherosclerosis in the ligated carotid artery was markedly increased in hypercholesterolemic *Mettl7a1^ΔECKO^* mice compared to hypercholesterolemic *Cdh5-Cre/ERT2* controls (n = 11 for *Cdh5-Cre/ERT2* mice; n =12 for *Mettl7a1^ΔECKO^* mice, *p* values were obtained using two-tailed Mann-Whitney U test using GraphPad Prism. *\*\*p* ≤ 0.01). Scale bar (100 mm) was shown in the right-lower corner of oil-red staining images. Data were presented as mean ± SD.

To establish the causal role of endothelial *Mettl7a1* in atherosclerosis, we generated a second mouse line in which *Mettl7a1* was selectively deleted in adult vascular endothelium (Supplemental Figure 6). Briefly, *Mettl7a1^fl/fl^* mice were engineered and then crossed with *Cdh5-Cre/ERT2* mice to generate *Cdh5-Cre/ERT2:Mettl7a1^fl/fl^ (Mettl7a1^ΔECKO^)* mice, in which *Mettl7a1* can be deleted in adult vascular endothelium by a tamoxifen-induced Cre recombinase driven by the endothelium-specific *Cdh5* (VE-Cadherin) promoter^36^. Carotid atherosclerosis in the ligated LCA was significantly increased in the PCSK9-AAV9 virus-injected *Mettl7a1^ΔECKO^* mice when compared to control *Cdh5-Cre/ERT2* mice subjected to tamoxifen and PCSK9-AAV9 viruses (Figure 2b), demonstrating that endothelial *Mettl7a1* deletion led to increased carotid atherosclerosis. These data collectively demonstrated that METTL7A is a previously unrecognized mechanosensitive methyltransferase governing the endothelial mRNA m^7^G methylome and endothelial METTL7A reduction causatively promotes atherogenesis *in vivo*.

### METTL7A expression is significantly reduced in human atherosclerosis

To assess whether METTL7A expression correlates with atherosclerotic plaques in humans, we analyzed RNA-seq data from ischemic atherosclerotic coronary artery tissue and non-atherosclerotic coronary arteries. Ischemic (n=35) and non-atherosclerotic (n=23) human coronary artery tissue specimens were obtained from diseased heart transplant donors^25^. Total RNA was extracted, subjected to quality control, and processed for RNA sequencing using the Illumina NovaSeq S4 Flowcell.

Differential gene expression analysis was conducted using DESeq2, comparing atherosclerotic and non-atherosclerotic arteries. As shown in Figure 3a, differential gene expression analysis revealed a significant reduction in *METTL7A* expression in ischemic atherosclerotic lesions compared to healthy controls. To further evaluate endothelial METTL7A expression in atherosclerotic lesions, immunostaining was performed on human coronary tissue with and without atherosclerosis. De-identified coronary biopsy samples were stained for CD31 (endothelial marker) and METTL7A, with Hematoxylin and Eosin staining used for pathological characterization of atherosclerosis. A trained pathologist classified samples as non-lesion areas, atherosclerotic lesions, or atherosclerotic lesions with calcification. Immunostaining quantification demonstrated that endothelial METTL7A in CD31+ cells was significantly reduced in atherosclerotic lesions compared to non-lesion arteries, with the lowest levels observed in calcified plaques (Figure 3b-c). These findings establish a correlation between reduced endothelial METTL7A expression and human coronary atherosclerosis, suggesting a potential role for METTL7A in endothelial dysfunction and plaque progression.

**Figure 3.**
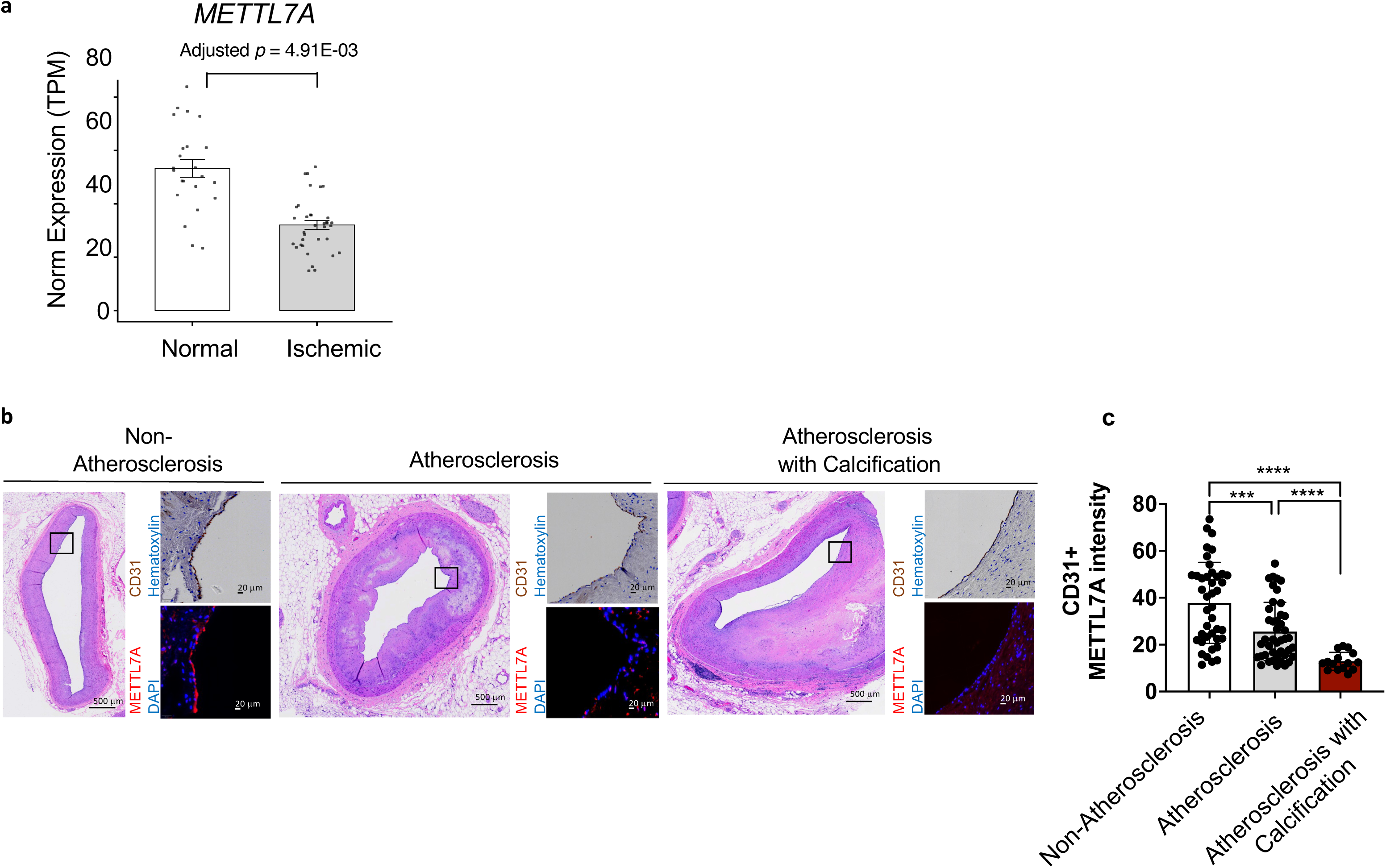
METTL7A expression is significantly reduced in human atherosclerosis. **(a)** RNA-seq analysis revealed significantly lower METTL7A mRNA expression in ischemic atherosclerotic coronary artery tissues (n = 35) compared to non-atherosclerotic coronary arteries (n = 23), Adjusted *p* = 4.91E-03. **(b–c)** Immunostaining of human coronary artery samples showed reduced METTL7A protein expression in CD31⁺ endothelial cells within atherosclerotic lesions, with the lowest levels observed in calcified plaques relative to non-lesion regions. Samples were pathologically classified as non-lesion, atherosclerotic lesion, or atherosclerotic lesion with calcification. Image counts: n = 39 (non-atherosclerotic controls), n = 43 (atherosclerotic lesions), n = 15 (calcified lesions). ***p ≤ 0.005, ****p ≤ 0.0001.

### METTL7A binds to protein-coding endothelial RNAs and mediates the flow-sensitive endothelial transcriptome

We next aimed to uncover vascular-related molecular mechanisms mediated by METTL7A. We determined the RNA substrates of METTL7A in an unbiased manner by conducting CLIP-seq (sequencing of *in vivo* protein-bound RNA targets by crosslinking and immunoprecipitation)^37^ in HAEC overexpressing HA-tagged METTL7A protein. RNA sequencing (RNA-seq) of the HA-pulled down METTL7A-RNA complex identified 2731 RNAs bound by endothelial METTL7A, 83.8% of which are in protein-coding mRNAs, 5.4% in long intergenic non-coding RNAs (lincRNA), 3.1% in microRNAs (miRNA), and 0.5% in ribosomal RNAs (rRNA) (Figure 4a). We identified U/A-AG-G/A as the top METTL7A RNA binding motif (Figure 4b), which was in agreement with previously identified m^7^G sites in tRNA (RAG<u>G</u>U)^14^ and mammalian mRNA (GA or GG-rich regions)^17,18^. Taken together, METTL7A CLIP-seq identified the RNA-binding motif of endothelial METTL7A and its preferential affinity to protein-coding mRNA. We then determined the flow-dependent, METTL7A-regulated endothelial transcriptome.

**Figure 4.**
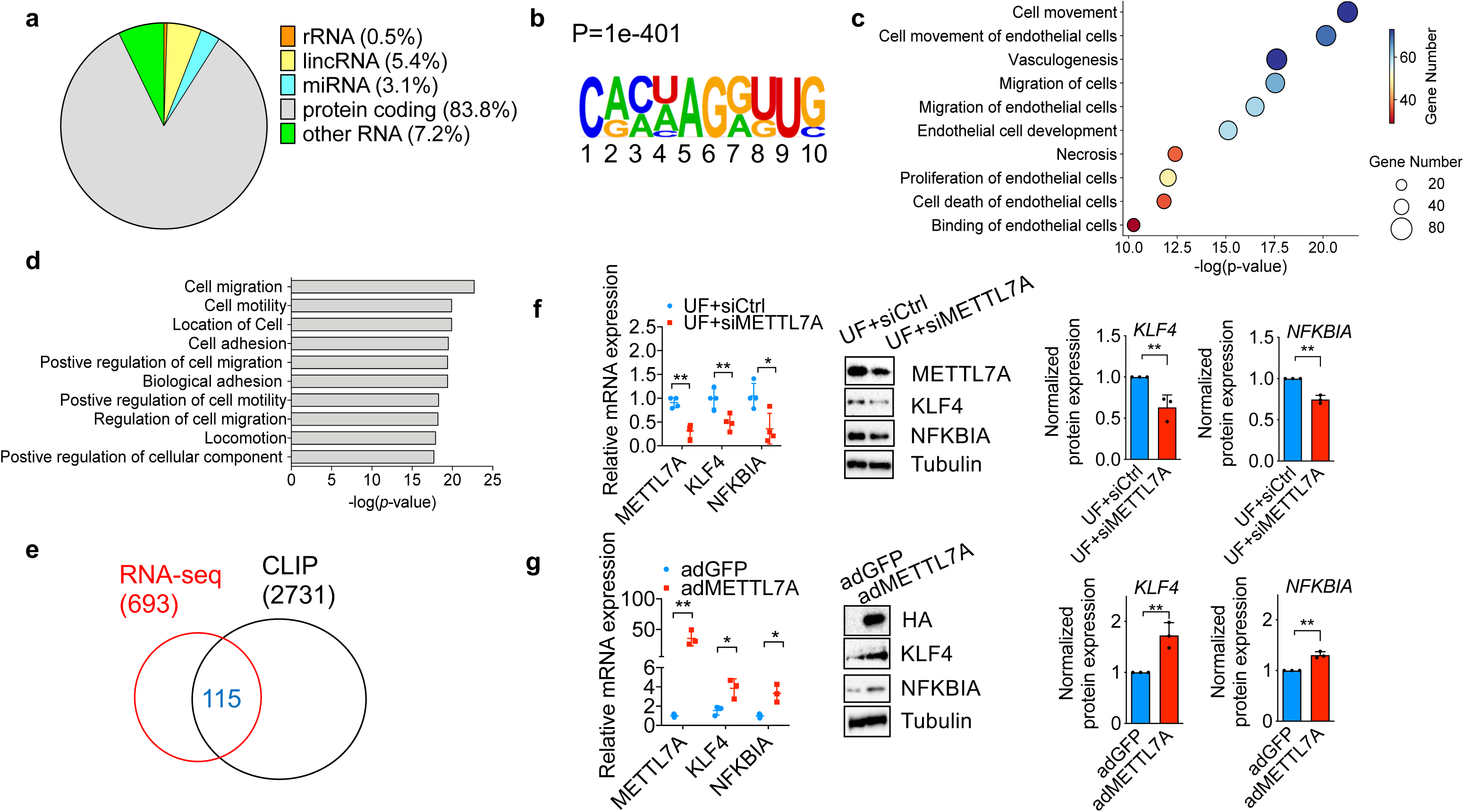
METTL7A binds to protein-coding endothelial RNAs and mediates the flow-sensitive endothelial transcriptome. **(a)** METTL7A CLIP-seq demonstrated that METTL7A preferentially binds to protein-coding mRNA in endothelial cells. **(b)** METTL7A CLIP-seq identified that U/A-AG-G/A is the top RNA binding motif of endothelial METTL7A. **(c)** Ingenuity Pathway Analysis (IPA) identified that endothelial cell movement, vasculogenesis, migration, and endothelial development are top annotated biological functions in the METTL7A-mediated endothelial transcriptome. **(d)** DAVID (the Database for Annotation, Visualization, and Integrated Discovery) identified that cell migration, cell motility, and cell adhesion are the top gene ontology (GO) biological processes regulated by endothelial METTL7A. **(e)** Comparison of the METTL7A CLIP-seq and METTL7A-mediated transcriptome identified a list of 115 endothelial mRNAs directly bound by METTL7A and significantly regulated by METTL7A. **(f)** METTL7A knockdown significantly reduced elevated mRNA and protein levels of KLF4 and NFKBIA in HAEC under unidirectional flow (n = 4 biological repeats; *p* values were obtained using two-tailed Student’s t-test using GraphPad Prism. \*\**p*≤0.01 and \**p*≤0.05). **(g)** Adenovirus-mediated METTL7A overexpression markedly increased the mRNA and protein expression of KLF4 and NFKBIA in HAEC (n = 3 biological repeats, \*\**p*≤0.01 and \**p*≤0.05). Data were presented as mean±SD.

Specifically, we conducted RNA-seq in HAEC under 24-hr unidirectional flow while the elevated METTL7A was suppressed by siRNA. RNA sequencing identified a total of 693 differentially expressed genes (DEGs) regulated by *METTL7A* knockdown, using a false discovery rate cutoff value of q < 0.1. Ingenuity Pathway Analysis (IPA)^29^ identified endothelial cell movement, vasculogenesis, migration, and endothelial development as top annotated biological functions in the METTL7A-regulated DEGs (Figure 4c).

Similarly, DAVID (the Database for Annotation, Visualization, and Integrated Discovery)^30^ analysis of the DEGs after *METTL7A* knockdown identified that cell migration, cell motility, and cell adhesion were among the top gene ontology (GO) biological processes (Figure 4d). Among the METTL7A-regulated DEGs, IPA bioinformatically identified a cohort of six transcriptional regulators with predicted widespread effects on the METTL7A-dependent transcriptome (Supplemental Table 2). Based on the direct METTL7A targets identified by CLIP-seq (total 2731 mRNAs) and the DEGs found from RNA-seq (total 693 mRNAs), we determined a list of 115 candidate METTL7A direct downstream targets (Figure 4e) (Supplemental Table 3).

From these identified METTL7A targets, we focused on Krüppel-like factor 4 (*KLF4*) and NFκB inhibitor alpha (*NFKBIA*). *KLF4* and *NFKBIA* were among the six IPA-predicted transcriptional regulators and were both downregulated by *METTL7A* knockdown demonstrated by RNA-seq. KLF4 is an evolutionarily conserved zinc finger-containing transcription factor that regulates diverse cellular processes such as cell migration, growth, proliferation, differentiation, apoptosis, and somatic cell reprogramming^38^. Moreover, NFKBIA encodes IκBα, a major inactivator of the NFκB-mediated signaling pathway, which binds to and inhibits the NFκB (nuclear factor-kappa-B) transcription factor^39^. Although how METTL7A might regulate KLF4 and NFKBIA remains unknown, the critical roles of *KLF4* and *NFKBIA* in endothelial health have been proposed^40–42^. *In vitro* and *in vivo* data demonstrated that UF upregulates *KLF4* expression and inactivates NFκB-mediated signaling to promote the anti-inflammatory, anti-adhesive, and anti-permeable endothelial phenotype resistant to atherosclerosis^34,40,43^. It has also been demonstrated that DF downregulates *KLF4* and activates NFκB-dependent pathways in inflamed endothelial cells prone to atherogenesis^27,44–46^. The significance of *KLF4* and *NFKBIA* in the *METTL7A*-mediated endothelial transcriptome is supported by the IPA analysis that *KLF4* and *NFKBIA* are highly connected hub genes involved in top regulated biological pathways of cell movement, cellular movement of endothelial cells, and migration of cell (Supplemental Figure 7). In addition, downregulation of *KLF4* and *NFKBIA* by *METTL7A* knockdown in HAEC under UF was validated by real-time PCR and Western Blots (Figure 4f).

Moreover, adenovirus-induced *METTL7A* overexpression significantly increased *KLF4* and *NFKBIA* expression in HAEC (Figure 4g). Taken together, *KLF4* and *NFKBIA* are METTL7A downstream targets and key regulatory hub genes governing the METTL7A-dependent endothelial transcriptome.

### METTL7A confers internal m^7^G modifications of *KLF4* and *NFKBIA* mRNAs and enhances their stability

To investigate whether METTL7A regulates endothelial *KLF4* and *NFKBIA* mRNA via internal m^7^G methylation, we performed RNA immunoprecipitation (RIP)-qPCR^17^. We used an anti-m^7^G-specific antibody and primer sets targeting a METTL7A binding site in *KLF4* or *NFKBIA* mRNA identified by METTL7A CLIP-seq. m^7^G-RIP-PCR demonstrated that in HAEC subjected to UF, METTL7A knockdown significantly reduced internal m^7^G in *KLF4* and *NFKBIA* mRNA (Figure 5a). Next, to support the role of METTL7A in mediating internal m^7^G methylation on *KLF4* and *NFKBIA*, we conducted an *in vitro* methylation assay (Figure 5b)^17,20,47^. We incubated recombinant METTL7A protein with *in vitro*-transcribed *KLF4* and *NFKBIA* mRNA molecules along with S-(5′-adenosyl)-L-methionine-d3 (d3-SAM) as the cofactor. The deuterium-substituted methyl group of d3-SAM allowed for precise methylation quantification by LC-MS/MS^17,20,47^, which showed that METTL7A effectively conferred internal m^7^G methylation on *KLF4* and *NFKBIA* transcripts *in vitro* (Figure 5b). Collectively, our data established METTL7A as an internal m^7^G methyltransferase for *KLF4* and *NFKBIA* mRNAs in HAEC.

**Figure 5.**
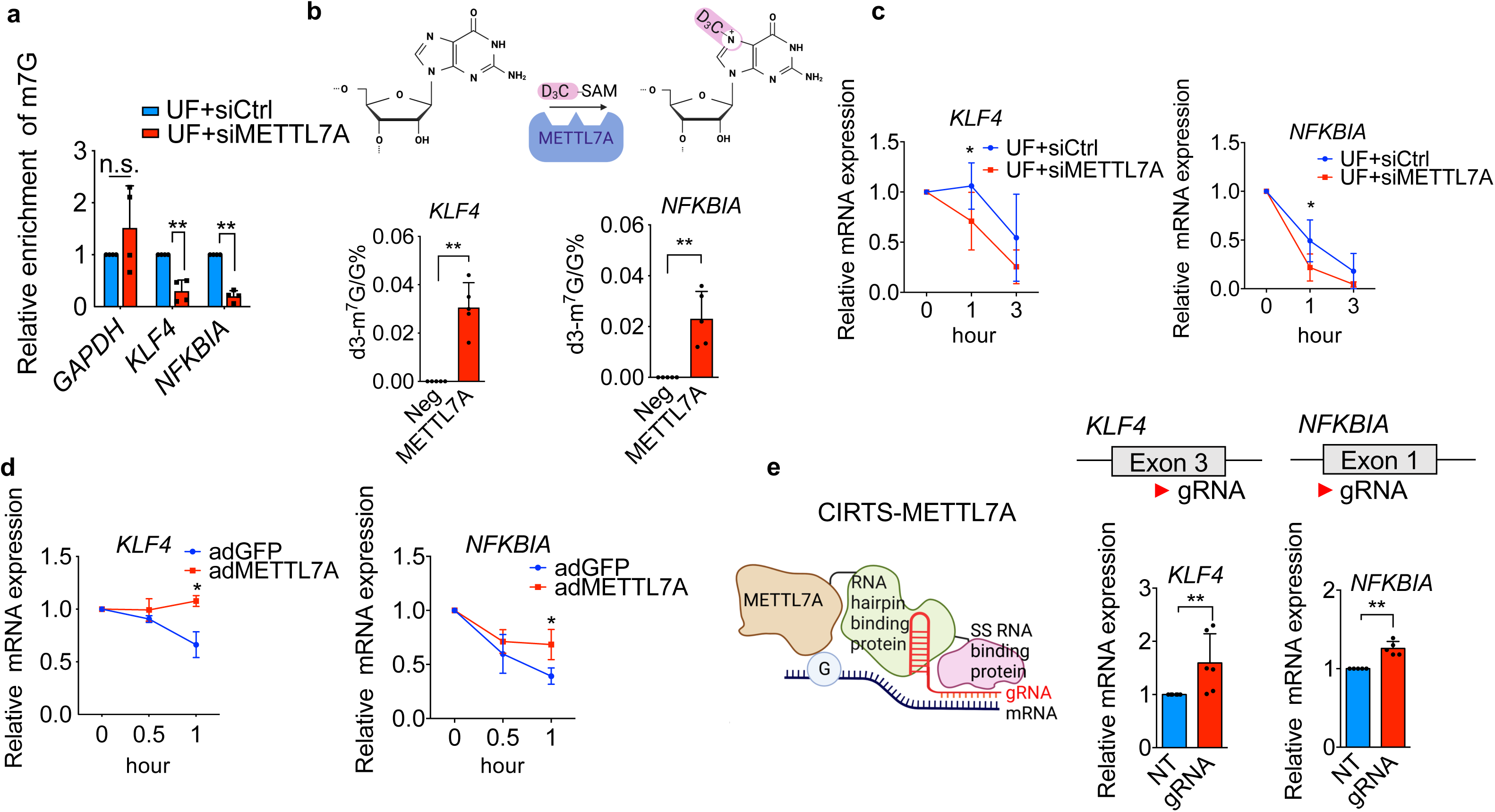
METTL7A functions as a methyltransferase to confer internal m^7^G modifications of *KLF4* and *NFKBIA* mRNAs and enhance their stability. (a) m^7^G-RIP-qPCR using anti-m^7^G-specific antibody and gene-specific primers demonstrated that METTL7A knockdown markedly reduced internal m^7^G in *KLF4* and *NFKBIA* mRNAs in HAEC. No change of internal m^7^G of endothelial *GAPDH* mRNA was detected (n = 4 with 2 biological repeats and 2 technical replicates, \*\**p*≤0.01 and n.s.= non-significance). *p* values were obtained using two-tailed Student’s t-test using GraphPad Prism. (b) *In vitro* methylation assays using purified METTL7A protein and S-(5′-Adenosyl)-L-methionine-d3 (d3-SAM) demonstrated that METTL7A effectively conferred internal m^7^G methylation on *KLF4* and *NFKBIA* transcripts (n = 5 independent reactions with *KLF4* transcript, n = 5 independent reactions with *NFKBIA* transcript, \*\**p*≤0.01). Negative control (Neg) is the experimental control without METTL7A protein. **(c)** Actinomycin D-based mRNA stability assays showed that knockdown of elevated METTL7A in HAEC under unidirectional flow markedly decreased the half-life of *KLF4* and *NFKBIA* transcripts (n = 4 biological repeats, \**p*≤0.05). **(d)** Adenovirus-mediated METTL7A overexpression markedly increased the stability of *KLF4* and *NFKBIA* transcripts in HAEC treated with Actinomycin D (n = 4 biological repeats, \**p*≤0.05). **(e)** Site-targeted delivery of METTL7A by CRISPR-Cas-inspired RNA targeting system (CIRTS) markedly increased *KLF4* and *NFKBIA* mRNA in HAEC. KLF4 or NFKBIA-targeting guide RNAs were designed against KLF4 or NFKBIA m^7^G sites validated by METTL7A CLIP-seq and m^7^G-RIP-PCR (n = 6 biological replicates for KLF4 targeting experiments and n = 5 biological replicates for NFKBIA targeting experiments, \*\**p*≤0.01). Data were presented as mean±SD.

Decreased *KLF4* and *NFKBIA* mRNA from *METTL7A* knockdown and increased *KLF4* and *NFKBIA* transcripts due to *METTL7A* overexpression suggested a mechanism that METTL7A enhances the RNA stability of its downstream targets. To test this hypothesis, we performed mRNA stability assays by treating cells with the transcription inhibitor actinomycin D and then tracked mRNA levels by qPCR at different time points. We showed that siRNA knockdown of UF-elevated METTL7A in HAEC significantly decreased the half-life of *KLF4* and *NFKBIA* transcripts (Figure 5c).

Concordantly, adenovirus-mediated *METTL7A* overexpression significantly increased the stability of *KLF4* and *NFKBIA* transcripts in actinomycin D-treated HAEC (Figure 5d). We further verified the causal relationship between METTL7A and its downstream targets *KLF4* and *NFKBIA* with a CRISPR-Cas-inspired RNA targeting system (CIRTS)^28^, which employs Watson-Crick-Franklin base pair interactions with a guide RNA (gRNA) to specifically deliver epitranscriptomic regulators to targeted transcripts. We engineered CIRTS to deliver METTL7A and designed guide RNAs targeting *KLF4* or *NFKBIA* m^7^G sites collectively identified by METTL7A CLIP-seq and m^7^G-RIP-PCR. CIRTS-METTL7A, along with *KLF4* or *NFKBIA*-targeting guide RNAs, markedly increased the abundance of *KLF4* and *NFKBIA* transcripts in HAEC compared to cells transfected with CIRTS-METTL7A and non-targeting guide RNAs (Figure 5e). Taken together, these results demonstrate that METTL7A functions as a previously unrecognized methyltransferase that confers internal m^7^G modifications on mRNAs to enhance their stability. In particular, endothelial METTL7A regulates expression and stability of *KLF4* and *NFKBIA,* key transcriptional regulators of vascular health.

### Endothelial METTL7A overexpression markedly increases *Klf4* and *Nfkbia* expression and lessens atherosclerosis *in vivo*

Through *in vivo* mouse studies, we further validated the critical role of endothelial METTL7A in enhancing *KLF4* and *NFKBIA* expression and reducing atherogenesis. In wild-type B6 mice, disturbed flow in the ligated LCA significantly reduced intimal (endothelial) expression of *Mettl7a1* (Figure 1e), accompanied by decreased *Klf4*, *Nfkbia*, compared to the RCA (Supplemental Figure 8). In addition, *Klf4* and *Nfkbia* transcripts were significantly reduced in isolated carotid (intimal) endothelial cells from *Mettl7a1^-/-^* mice compared to those isolated from wild-type mice (Figure 6a). In hypercholesterolemic (PCSK9 virus-injected) *Mettl7a1^-/-^*mice in which atherosclerosis was markedly increased in the ligated carotid artery, immunostaining demonstrated that *Klf4* and *Nfkbia* expression was significantly reduced compared to hypercholesterolemic *Mettl7a1^+/+^* mice (Figure 6b). We verified the causal role of endothelial *METTL7A* in *Klf4* and *Nfkbia* expression and in athero-protection *in vivo* through nanoparticle-mediated endothelial *METTL7A* restoration in *Mettl7a1^-/-^* mice (Figure 6c). We employed cationic polymer-based nanoparticles to deliver *METTL7A*-expressing plasmids driven by an endothelial-specific VE-cadherin/*CDH5* promoter, an approach that effectively overexpresses the gene of interest in adult vascular endothelium *in vivo*^48,49^.

**Figure 6.**
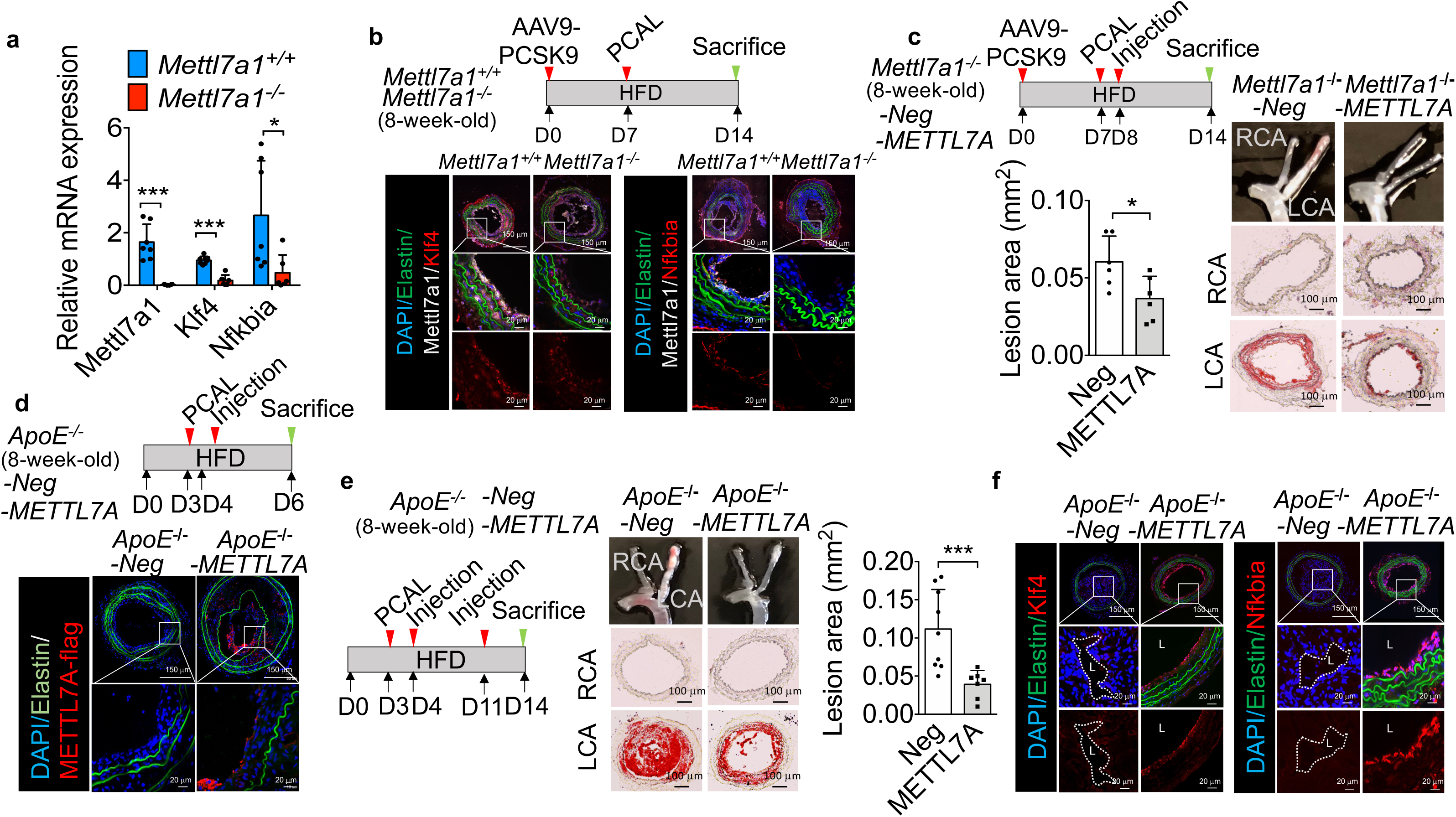
Restoration of endothelial METTL7A using CDH5 promoter-driven plasmids delivered via polymer-based nanoparticles significantly upregulates Klf4 and Nfkbia expression and reduces disturbed flow-induced atherosclerosis *in vivo*. (a) Endothelial Klf4 and Nfkbia transcripts were significantly reduced in carotid intima in *Mettl7a1^-/-^* mice compared to *Mettl7a1^+/+^* mice (*Mettl7a1^+/+^* n = 7 individual mice, *Mettl7a1^-/-^* n = 6 individual mice, \*\*\**p*≤0.001 and \**p*≤0.05). *p* values were obtained using two-tailed Student’s t-test using GraphPad Prism. **(b)** Klf4 and Nfkbia protein expression was significantly reduced in the ligated carotid artery in hypercholesterolemic (PCSK-9 virus-injected) *Mettl7a1^-/-^* mice compared to hypercholesterolemic *Mettl7a1^+/+^* mice. DAPI shows blue color; elastin shows green color; Mettl7a1 shows white color and Klf4 or Nfkbia shows red color in the images. Scale bars (150 and 20 μm) were indicated in the right-lower corner of immunostaining images. The zoomed-in area is 150 μm x 150 μm. **(c)** Atherosclerosis in the ligated carotid artery in hypercholesterolemic *Mettl7a1^-/-^* mice is markedly lessened by an injection of nanoparticles encapsulating METTL7A-expressing plasmids driven by an endothelial-specific VE-cadherin/CDH5 promoter, when compared to lesions in hypercholesterolemic *Mettl7a1^-/-^* mice injected with nanoparticles carrying control plasmids with CDH5 promoter but no expressed gene (Neg). (n = 6 individual mice, \**p*≤0.05). Scale bar (100 μm) was shown in the right-lower corner of oil-red staining images. **(d)** An injection of CDH5-METTL7A plasmid-encapsulated nanoparticles effectively induced intimal expression of METTL7A in the ligated carotid artery of hypercholesterolemic *ApoE^−/−^* mice. DAPI shows blue color; elastin shows green color and FLAG-tagged METTL7A shows red color in the images. Scale bars (150 and 20 μm) were indicated in the right-lower corner of immunostaining images. The zoomed-in area is 150 μm x 150 μm. **(e)** Atherosclerosis in the ligated carotid artery in hypercholesterolemic *ApoE^-/-^* mice is markedly lessened by injections of nanoparticles encapsulating CDH5-METTL7A plasmids, when compared to lesions in hypercholesterolemic in *ApoE^-/-^* mice injected with nanoparticles carrying control plasmid (n = 9 individual mice for negative control and 7 individual mice for METTL7A overexpression condition, \*\*\**p*≤0.005). *p* values were obtained using two-tailed Mann-Whitney U test using GraphPad Prism. Scale bar (100 μm) was shown in the right-lower corner of oil-red staining images. **(f)** Intimal protein expression Klf4 and Nfkbia in the ligated carotid artery was increased in hypercholesterolemic *ApoE^-/-^* mice administered with nanoparticles encapsulating CDH5-METTL7A plasmids, when compared to hypercholesterolemic *ApoE^-/-^* mice subjected to nanoparticles carrying control plasmids. L indicates lumen area. DAPI shows blue color; elastin shows green color and klf4 or nfkbia shows red color in the images. Scale bars (150 and 20 μm) were shown in the right-lower corner of immunostaining images. The zoomed-in area is 150 μm x 150 μm.

Nanoparticles encapsulating *CDH5-METTL7A* plasmid successfully increased METTL7A expression in the endothelium but not in the underlying media and adventitia in *Mettl7a1^-/-^* mice, demonstrated by real-time qPCR (Supplemental Figure 9).

Endothelial *METTL7A* restoration via nanoparticle delivery markedly reduced DF-induced atherosclerosis in the ligated left carotid artery (LCA) of *Mettl7a1^-/-^* mice (Figure 6c). These findings further support the causal role of mechanosensitive *Mettl7a1* in regulating endothelial *Klf4* and *Nfkbia* expression and highlight the therapeutic potential of restoring endothelial METTL7A to mitigate DF-induced vascular complications.

To further establish that endothelial *METTL7A* overexpression is an attractive approach in lessening atherosclerosis, we employed nanoparticle-mediated endothelial *METTL7A* overexpression in *ApoE^−/−^* mice fed a high-fat diet. Similar to the results in wild-type B6 mice, we detected a significant reduction of endothelial *Mettl7a1* in the ligated LCA exposed to DF in *ApoE^−/−^*mice (Supplemental Figure 10). An injection of nanoparticles encapsulating the *CDH5-METTL7A* plasmid effectively induced intimal (endothelial) expression of *METTL7A* in the ligated LCA of hypercholesterolemic *ApoE^−/−^* mice (Figure 6d). We then tested the therapeutic effectiveness of injections of *CDH5-METTL7A* plasmid-encapsulated nanoparticles in reducing DF-induced atherosclerosis. Carotid atherosclerosis in the ligated LCA in *ApoE^−/−^* mice was significantly reduced by endothelial overexpression of *METTL7A* (Figure 6e) and was accompanied by elevated endothelial expression of *Klf4* and *Nfkbia* (Figure 6f).

### Nanomedicine-mediated targeted delivery of METTL7A mRNA to inflamed endothelial cells attenuates atherosclerosis *in vivo*

We next tested the hypothesis that restoration of METTL7A in inflamed endothelial cells—such as those activated by local disturbed blood flow—could mitigate atherogenesis. We reasoned that targeted delivery of METTL7A mRNA to endothelial cells expressing vascular cell adhesion molecule-1 (VCAM-1) would be an ideal strategy, given that VCAM-1 is minimally expressed in quiescent endothelium but significantly upregulated under disturbed flow^50,51^. To accomplish this, we employed a VCAM1-targeting nanoparticle system recently developed in our lab, which efficiently delivers therapeutic mRNA to VCAM1-expressing endothelial cells *in vivo*^33^. The self-assembled lipid nanoparticles were composed of dioleoylphosphatidylethanolamine (DOPE), lipid-grafted polyamidoamine dendrimer (G0-C14), cholesterol, 1,2-distearoyl-sn-glycero-3-phosphoethanolamine-poly(ethylene glycol) (DSPE-PEG), and mRNA. To enable endothelial targeting, the VCAM1-binding peptide VHPKQHR^52^, identified via phage display, was conjugated to DSPE-PEG, resulting in its surface presentation on the nanoparticle corona. Transmission electron microscopy (TEM) revealed monodisperse nanoparticles with a dry diameter of ∼70 nm and a hydrodynamic diameter of ∼120 nm in deionized water^33^.

Chemical modifications of mRNA have been shown to improve translational efficiency and reduce immunogenicity^53–55^. Therefore, METTL7A mRNA was chemically modified with N1-methylpseudouridine (m^1^Ψ). Nanoparticles encapsulating a non-translatable mutant METTL7A mRNA served as negative controls. We first evaluated the therapeutic efficacy of these nanoparticles in *Mettl7a1^-/-^*mice subjected to AAV9-PCSK9 injection, high-fat diet, and partial carotid ligation. A single intravenous injection of METTL7A m1Ψ mRNA-loaded VCAM1-targeting nanoparticles significantly reduced disturbed flow-induced carotid atherosclerosis in the ligated artery, compared to mice treated with PBS or control nanoparticles (Figure 7a). We further validated the therapeutic effect in *ApoE^−/−^* mice subjected to a high-fat diet and partial carotid ligation. Two intravenous injections of METTL7A mRNA-encapsulated, VCAM1-targeting nanoparticles markedly attenuated lesion formation, relative to control groups (Figure 7b). Consistent with reduced atherosclerosis, endothelial expression of *Klf4* and *Nfkbia* was significantly upregulated in the ligated carotid arteries of METTL7A mRNA-treated mice (Figure 7c). These results not only further support the atheroprotective role of endothelial METTL7A but also provide proof of concept for a targeted, nanomedicine-based mRNA therapeutic strategy to restore METTL7A expression and attenuate atherosclerosis.

**Figure 7.**
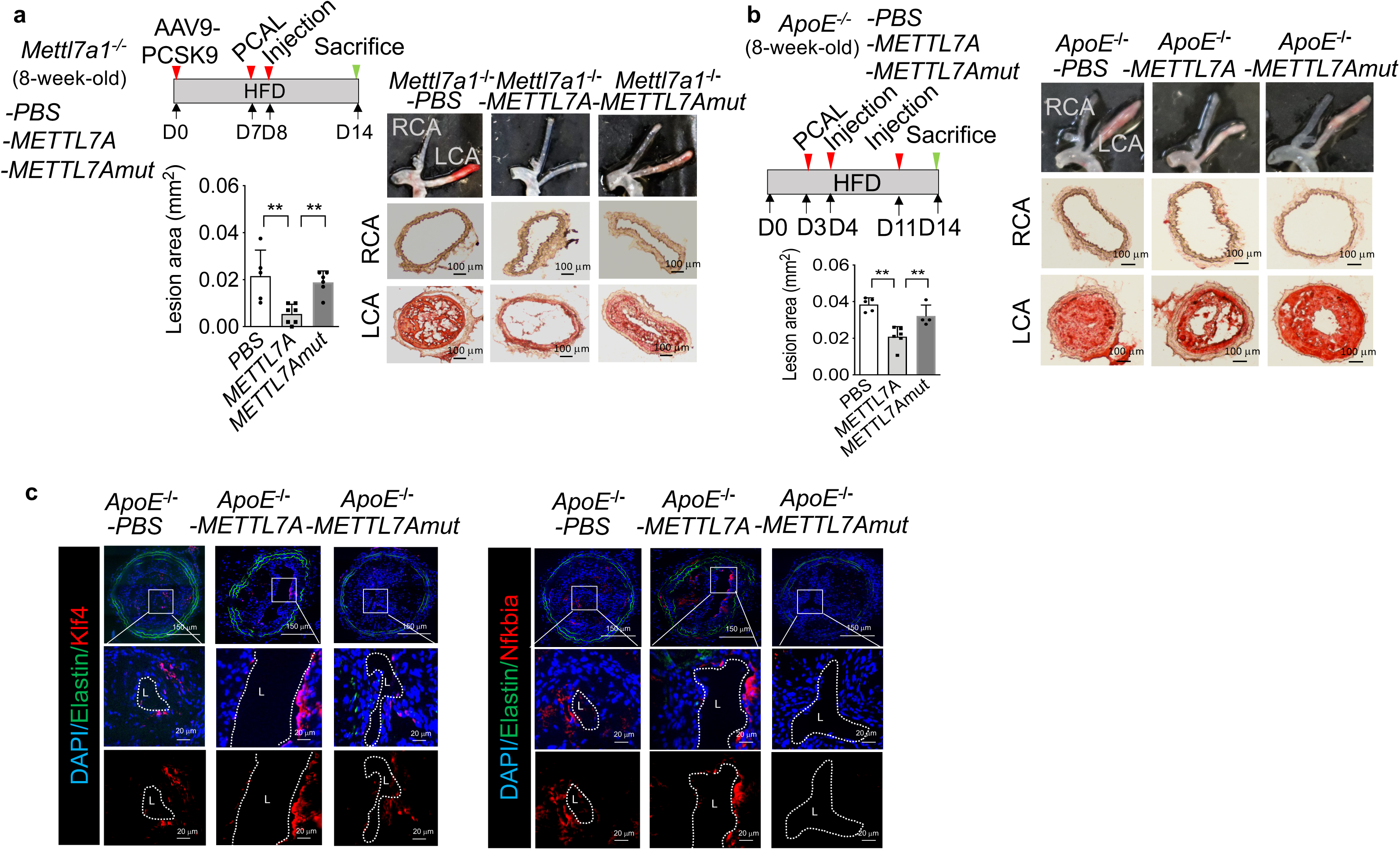
Restoration of METTL7A in inflamed endothelial cells using N1-methylpseudouridine (m1Ψ)-modified METTL7A mRNA delivered via VCAM1-targeting lipid nanoparticles attenuates atherosclerosis *in vivo*. **(a)** In hypercholesterolemic *Mettl7a1^-/-^* mice, a single intravenous injection of VCAM1-targeting lipid nanoparticles encapsulating m1Ψ-modified METTL7A mRNA (METTL7A) significantly reduced disturbed flow-induced atherosclerosis in the ligated carotid artery compared to mice treated with PBS or nanoparticles carrying non-translatable mutant METTL7A mRNA (METTL7Amut) (n = 5 for PBS, n = 7 for METTL7A mRNA, n = 5 for METTL7Amut; ** p ≤ 0.01). Scale bar: 100 μm. **(b)** In hypercholesterolemic *ApoE^-/-^* mice, two intravenous injections of VCAM1-targeting lipid nanoparticles encapsulating m1Ψ-modified METTL7A mRNA significantly attenuated atherosclerotic lesion formation in the ligated carotid artery compared to PBS or control nanoparticle groups (n = 5 for PBS, n = 6 for METTL7A mRNA, n = 4 for METTL7Amut; ** p ≤ 0.01). Scale bar: 100 μm. Statistical analyses were performed using a two-tailed Mann– Whitney U test (GraphPad Prism). **(c)** Immunostaining demonstrated increased intimal expression of KLF4 and NFKBIA proteins in the ligated carotid arteries of *ApoE^-/-^* mice treated with METTL7A mRNA nanoparticles compared to METTL7Amut. Lumen (L) is indicated. Nuclei (DAPI) are shown in blue, elastin in green, and KLF4 or NFKBIA in red. Scale bars: 150 μm (overview) and 20 μm (zoomed-in panels). Zoomed-in regions measure 150 μm × 150 μm.

## Discussion

In summary, we identify internal m^7^G methylation of mRNA as a previously unrecognized molecular regulator of mechanotransduction—a fundamental biological process essential for maintaining tissue homeostasis^56–59^, particularly in the endothelium, where unidirectional flow promotes vascular quiescence, while disturbed flow drives endothelial dysfunction^2–6^. Aberrant cellular responses to mechanical forces contribute to the development of numerous human diseases, including those affecting the cardiovascular, pulmonary, orthopedic, muscular, and reproductive systems^57^. Our findings demonstrate that METTL7A governs the mechanosensitive internal m^7^G methylome of endothelial mRNA, enhancing the stability of transcripts of key anti-inflammatory genes and supporting vascular health. Furthermore, targeted restoration of endothelial METTL7A offers a promising therapeutic strategy for vascular wall–based interventions aimed at counteracting disturbed flow-induced complications such as atherosclerosis.

Although m^7^G is well established for its essential roles at the 5′ cap of mRNA^11^ and within non-coding RNAs such as tRNA^14^ and rRNA^13^, internal m^7^G methylation in mammalian mRNA has only recently been discovered^17^, expanding its potential roles in post-transcriptional gene regulation. *Our in vivo* and *in vitro* data reveal that internal m^7^G methylation in endothelial mRNA is dynamically regulated by biomechanical forces through the mechanosensitive methyltransferase METTL7A. By mapping genome-wide METTL7A RNA-binding sites and motifs, and analyzing transcriptomic changes following METTL7A perturbation, we identified KLF4 and NFKBIA as direct downstream targets—highlighting a novel epitranscriptomic mechanism by which endothelial cells convert mechanical cues into protective gene expression programs. Given the widespread expression of KLF4 and NFKBIA in various tissues and their major roles in key cellular functions such as inflammation, proliferation, differentiation, and somatic cell reprogramming^38,60^, METTL7A may participate in a variety of biological processes in addition to vascular health. For example, single-cell indexing lineage mapping in cultured cells suggested that Mettl7a1 functions as a pro-reprogramming factor to stabilize induced endoderm progenitor reprogramming from fibroblasts although the underlying molecular action remains unclear^61^.

We established the role of METTL7A in mediating internal m^7^G methylation in the context of the vascular system. With an *in vitro* methylation assay, we validated that METTL7A adds internal m^7^G on KLF4 and NFKBIA transcripts. CIRTS-METTL7A^28^ and RNA stability assays corroborated METTL7A-mediated increase of *KLF4* and *NFKBIA* transcripts and illuminated that the METTL7A-m^7^G axis controls the endothelial transcriptome by enhancing stability of direct mRNA targets. In non-endothelial cell lines (HeLa, HEK293T, A549, or Caco-2 cells) where internal m^7^G of mammalian mRNA and miRNA was first discovered, METTL1-mediated m^7^G was shown to increase mRNA translation efficiency or augment the miRNA maturation^16,17^. Recently, it was discovered that Quaking proteins (QKIs) bind to mRNAs with internal m^7^G modifications at GA-rich motifs, leading to an increase in the overall mRNA half-life^18^. Our results along with previous data provide evidence that internal m^7^G could impact mRNA function and metabolism by a wide range of molecular actions. Despite the growing recognition of internal m^7^G as a regulatory epitranscriptomic mark, the molecular identity of its reader proteins and the mechanistic basis by which they recognize and regulate internal m^7^G– modified transcripts—particularly in the context of vascular biology and cardiovascular disease—remain largely undefined. Elucidating these mechanisms will be essential for uncovering how m^7^G methylation integrates into broader post-transcriptional regulatory networks governing endothelial function and vascular homeostasis.

Our new animal models and nanomedicine approaches demonstrated that endothelial METTL7A confers internal m^7^G methylation of mRNAs and lessens atherogenesis. Previously, the role of METTL7A and mRNA m^7^G in cardiovascular complication *in vivo* was virtually unknown. Here we showed that mice with global or endothelial-specific deletion of *Mettl7a1* significantly exacerbated atherosclerosis. In agreement with the *in vitro* results, *Mettl7a1* deletion *in vivo* remarkedly reduced endothelial internal m^7^G within endothelial mRNA and suppressed *Klf4* and *Nfkbia* expression.

Consistent with the findings in mouse models demonstrating the athero-protective role of endothelial METTL7A, RNA-seq and immunostaining analyses of human coronary arteries revealed a significant reduction in endothelial METTL7A expression within atherosclerotic lesions compared to non-lesion areas. These results highlight the clinical relevance of endothelial downregulation of mechano-sensitive METTL7A in vascular dysfunction and underscore its potential as a therapeutic target for vascular wall-based treatment of atherosclerosis. While the role of endothelial METTL7A loss in other vascular complications remains to be fully elucidated, several lines of evidence suggest that suppression of this mechanosensitive regulator may contribute more broadly to cardiovascular pathologies. Many vascular disorders— including peripheral artery disease, vascular aneurysms, arteriovenous fistula failure, aortic valvular disease, and graft failure—predominantly arise in regions of disturbed blood flow, where endothelial dysfunction is a central pathological feature^62^. Notably, reduced expression of endothelial KLF4 and/or NFKBIA has been implicated in pathological vascular remodeling and dysfunction across various disease contexts^63–68^. Our findings suggest that restoring endothelial METTL7A expression may offer a novel molecular mechanism to enhance the stability of mRNAs encoding transcriptional regulators, such as KLF4 and NFKBIA, whose transcripts are known to exhibit rapid decay compared to other functional gene classes^69,70^. This stabilization mechanism may extend to additional regulatory targets and provide a broader foundation for vascular gene therapy.

In addition to the internal m⁷G modification characterized in this study, it is likely that multiple epitranscriptomic regulators act in concert to modulate the mechano-sensitive endothelial transcriptome. For example, METTL3-mediated N⁶-methyladenosine (m⁶A) has been implicated in regulating genes critical for endothelial homeostasis. Disturbed flow has been shown to upregulate METTL3 and result in m⁶A hypermethylation of specific transcripts, leading to endothelial inflammation^71^ and glycolytic reprogramming^72^. However, the dynamic regulation of endothelial METTL3 by flow and its precise contribution to atherogenesis remain incompletely understood and warrant further investigation^71–73^.

To therapeutically exploit the vascular-protective function of endothelial METTL7A, we developed and validated two complementary precision nanomedicine strategies for targeted endothelial delivery of functional METTL7A. A polymer-based nanoparticle system was employed to deliver CDH5 promoter-driven METTL7A plasmids, while a lipid-based nanoparticle platform was used to deliver N1-methylpseudouridine (m^1^Ψ)-modified METTL7A mRNA specifically to inflamed endothelial cells. Both approaches significantly attenuated disturbed flow–induced atherosclerosis in mouse models. These findings establish proof of concept that therapeutic restoration of endothelial METTL7A, and by extension, targeted modulation of internal mRNA m^7^G methylation, represents a promising strategy against pathological vascular remodeling. Given the central role of endothelial dysfunction in vascular disease, these results position METTL7A-directed epitranscriptomic therapy as a novel and potentially transformative approach to combat atherosclerosis, the leading cause of morbidity and mortality worldwide^1^.

## Acknowledgments

This work was supported by the National Institutes of Health (NIH) grants HL161244 and HL155909, the American Heart Association (AHA) grant 23EIA1038679 and 24POST1198682, and the Gracias Family Foundation.

---

**Supplemental Figure 1.**
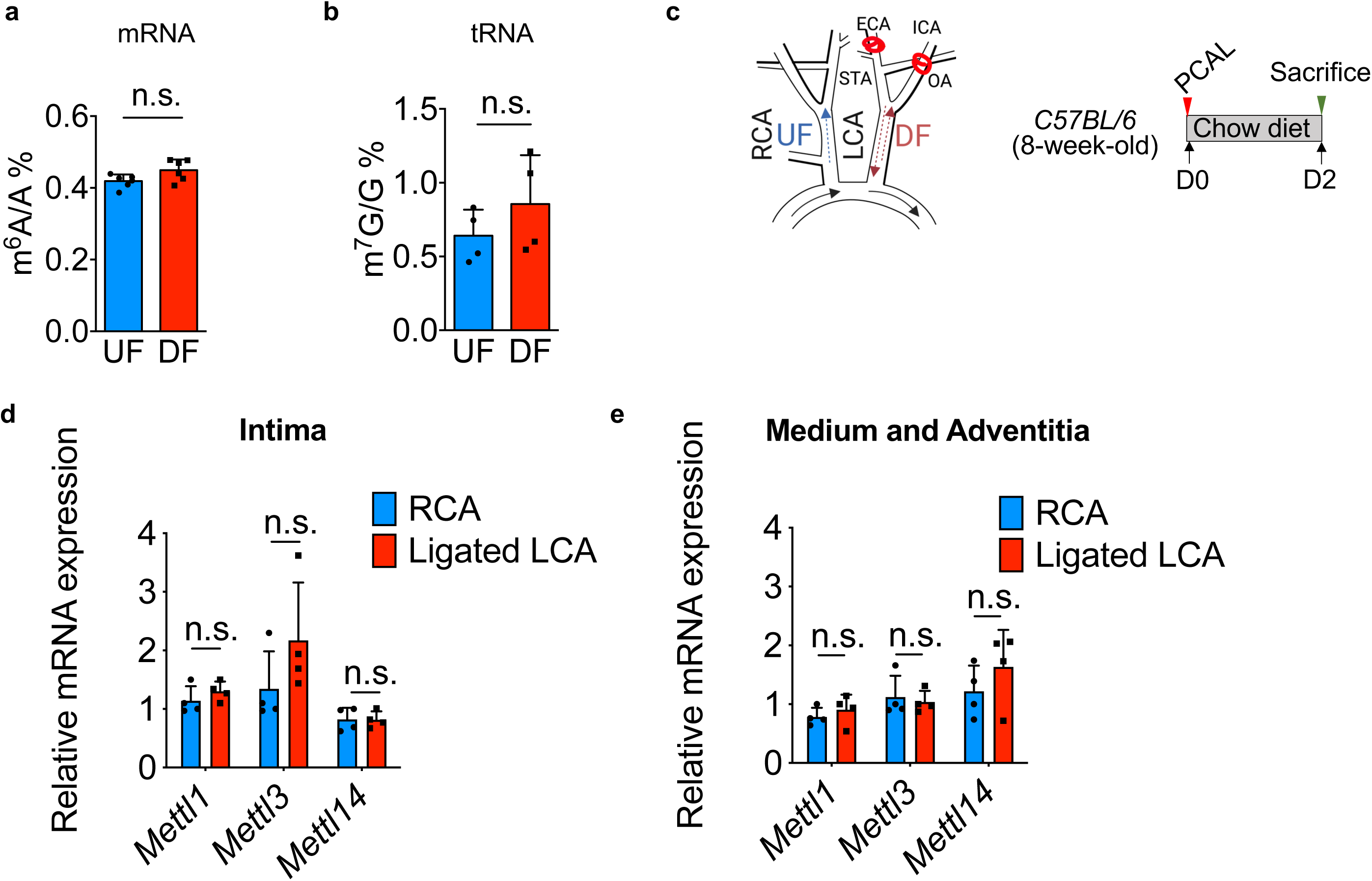
Internal m^6^A of endothelial mRNA and total m^7^G of endothelial tRNA are not regulated by atherosclerosis-relevant hemodynamic forces. Vascular expression of Mettl1, Mettl3, and Mettl14 is not regulated by acute disturbed flow *in vivo*. LC-MS/MS demonstrated that in HAEC subjected to 24-hr athero-protective unidirectional flow (UF) and athero-prone disturbed flow (DF), levels of **(a)** Internal m^6^A of mRNA (n = 6 biological repeats) and **(b)** m^7^G of tRNA (n = 4 biological replicates) were not changed. **(c)** Partial carotid ligation (PCAL) was performed to induce disturbed flow (DF) in *C57BL/6* mouse left carotid artery (LCA). **(d and e)** Real-time PCR demonstrated that 48-hr acute disturbed flow in the partially-ligated LCA had no effect on mRNA levels of *Mettl1*, *Mettl3*, and *Mettl14* in the mouse carotid intima and carotid media/adventitia when compared to those in the non-ligated right carotid artery (RCA) (n = 4 biological repeats). *p* values were obtained using two-tailed Student’s t-test using GraphPad Prism. n.s. indicates non-significance (*p*>0.05). Data were presented as mean±SD.

**Supplemental Figure 2.**
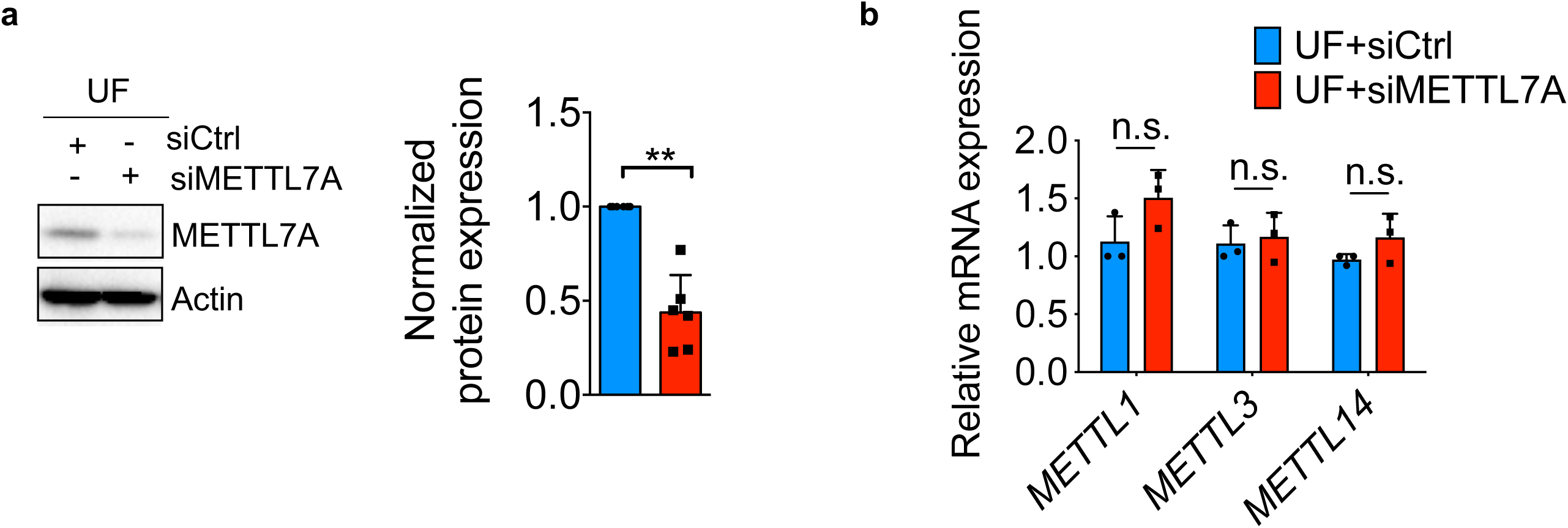
METTL7A knockdown has no effect on the endothelial expression of METTL1, METTL3, and METTL14. **(a)** METTL7A-targeting siRNA significantly decreased the METTL7A protein expression in HAEC under unidirectional flow (n = 6 biological repeats, \*\**p*≤0.01). **(b)** METTL7A knockdown had no effect on the expression of *METTL1*, *METTL3*, and *METTL14* in HAEC under unidirectional flow (n = 3 biological repeats). *p* values were obtained using two-tailed Student’s t-test using GraphPad Prism. n.s. indicates non-significance (*p*>0.05). Data were presented as mean±SD.

**Supplemental Figure 3.**
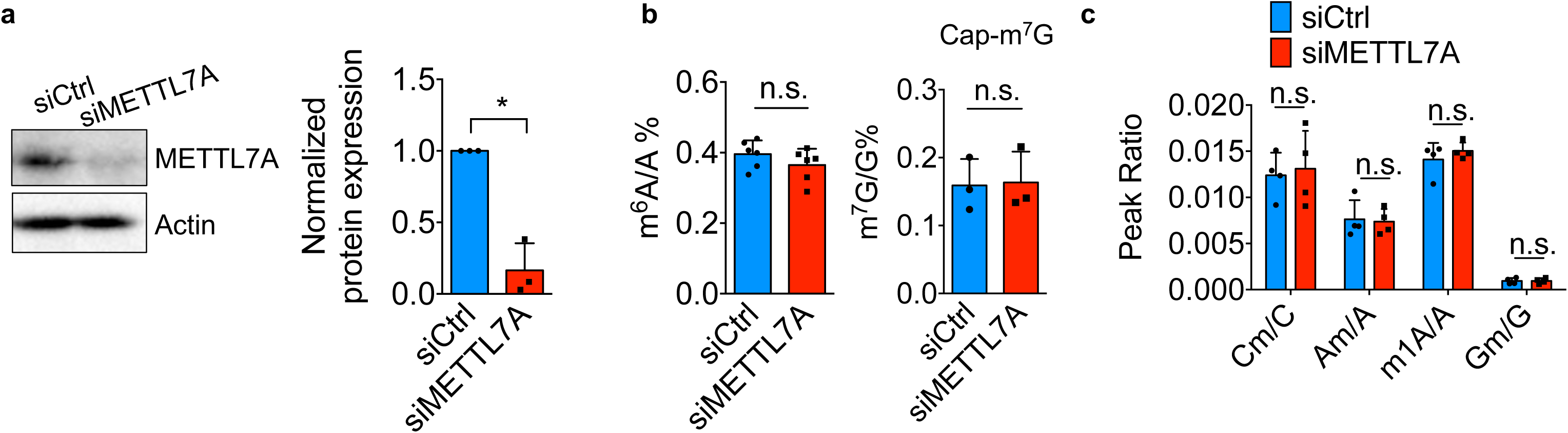
METTL7A knockdown has no effect on m^6^A/A, C^m^/C, A^m^/A, m^1^A/A and G^m^/G of endothelial mRNA. **(a)** METTL7A-targeting siRNA significantly decreased the METTL7A protein expression in HAEC (n = 3 biological repeats, \**p*≤0.05). **(b)** METTL7A knockdown had no effect on m^6^A and cap-m^7^G of mRNA in HAEC (n = 6 biological repeats for m^6^A measurement and 3 biological repeats for cap-m^7^G measurement). **(c)** METTL7A knockdown had no effect on C^m^/C, A^m^/A, m^1^A/A and G^m^/G of mRNA in HAEC (n = 4). *p* values were obtained using two-tailed Student’s t-test using GraphPad Prism. n.s. indicates non-significance (*p*>0.05). Data were presented as mean±SD.

**Supplemental Figure 4.**
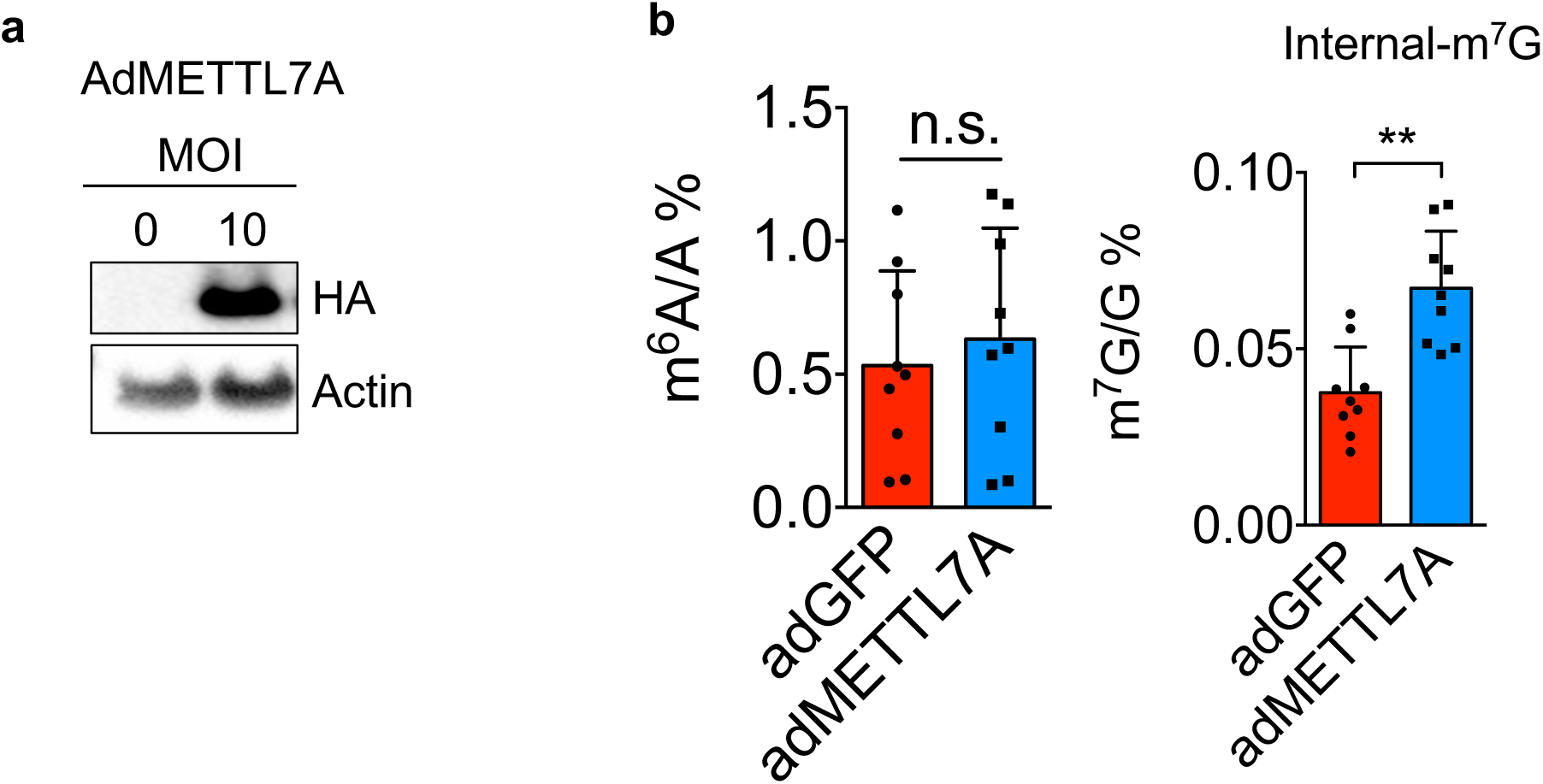
METTL7A overexpression significantly increases internal m^7^G but not the m^6^A of endothelial mRNA. **(a)** Adenoviruses (10 MOI/multiplicity of infection) effectively overexpressed HA-tagged METTL7A in HAEC as shown in the western blot. **(b)** Adenovirus-mediated METTL7A overexpression significantly increased the internal m^7^G but not the m^6^A of endothelial mRNA in HAEC (n = 9 biological repeats, \*\**p*≤0.01 and n.s. indicates non-significance (*p*>0.05)). Data were presented as mean±SD.

**Supplemental Figure 5.**
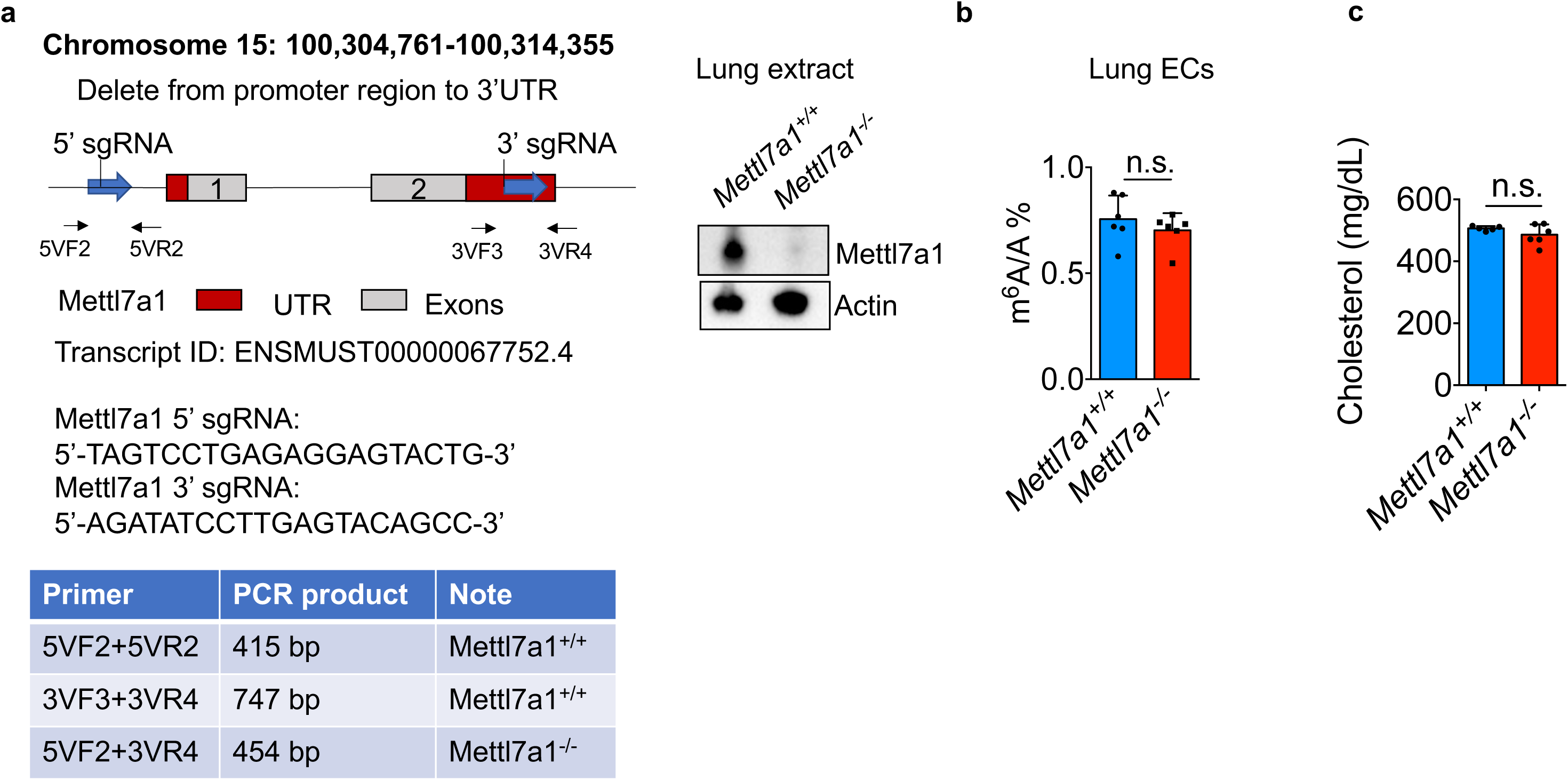
Mettl7a1 knockout in mice has no effect on the internal m^6^A of endothelial mRNA and cholesterol level. **(a)** *Mettl7a1^-/-^ C57BL/6* mice were generated by CRISPR Cas9-mediated gene deletion using a pair of guide RNAs (sgRNA) targeting the promoter region and downstream of exon 2 of Mettl7a1, validated by markedly reduced Mettl7a1 protein in lung tissues of *Mettl7a1^-/-^* mice compared to those in the wild-type *Mettl7a1^+/+^* mice. The genotyping primers (5VF2+3VR4) that were used to identify Mettl7a1 knockout are shown in the table below. **(b)** LC-MS/MS detected no change of internal m^6^A of mRNA in isolated lung endothelial cells from *Mettl7a1^-/-^* mice when compared to those in endothelial cells isolated from *Mettl7a1^+/+^* mice (n = 6 biological repeats). **(c)** Plasma cholesterol level was not affected in *Mettl7a1^-/-^* mice fed a high-fat diet compared to *Mettl7a1^-/-^* mice fed a high-fat diet (n = 5 for *Mettl7a1^+/+^* mice, n = 6 for *Mettl7a1^-/-^* mice). *p* values were obtained using two-tailed Student’s t-test using GraphPad Prism. n.s. indicates non-significance (*p*>0.05). Data were presented as mean±SD.

**Supplemental Figure 6.**
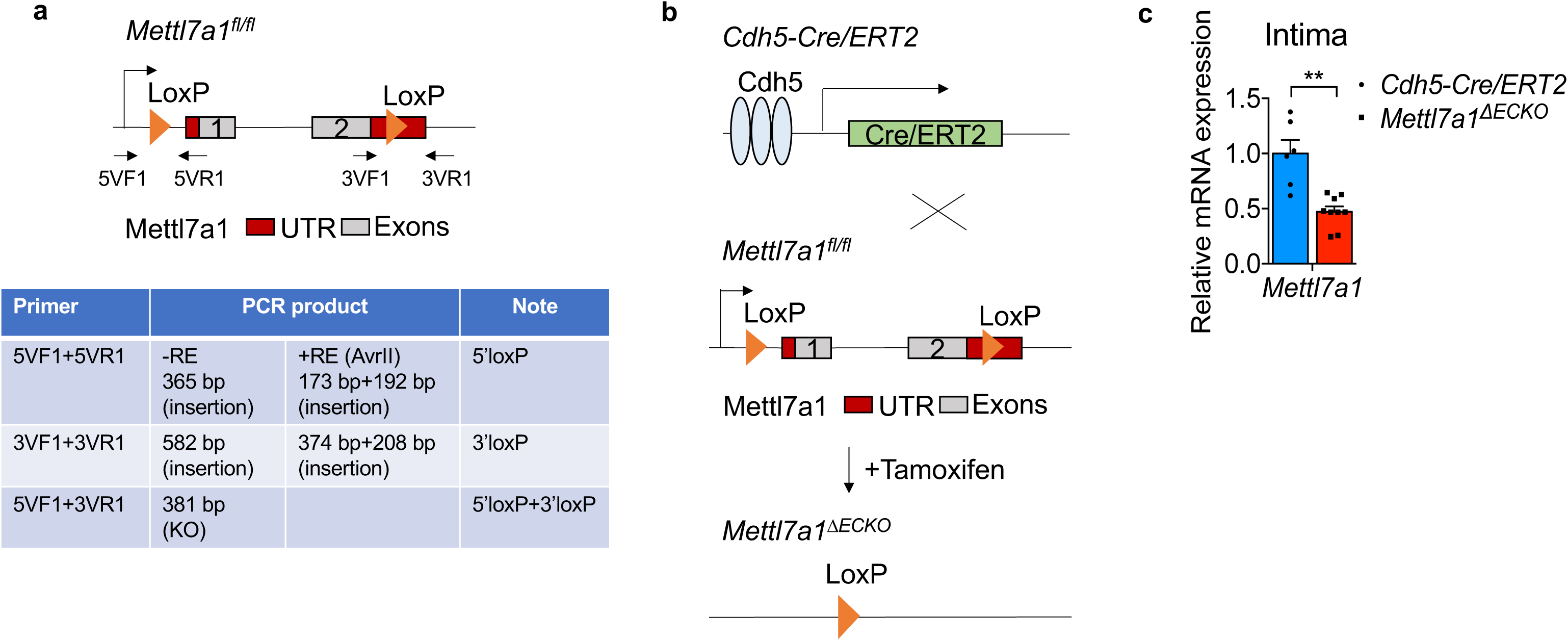
Endothelial Mettl7a1 expression is significantly reduced in *Mettl7a1^ΔECKO^* mice. **(a)** *Mettl7a1 ^fl/fl^ C57BL/6* mice were generated by CRISPR Cas9-mediated gene deletion using a pair of guide RNAs (sgRNA) and two single strand donor oligodeoxynucleotides (ssONDs), which are listed in the methods. The genotyping primers (5VF1+3VR1) that were used to identify 5’-loxP and 3’-loxP sites are listed in the table below. **(b)** *Cdh5-Cre/ERT2::Mettll7a1^fl/fl^* (*Mettl7a1^ΔECKO^*) mice were engineered to specifically knockout Mettl7a1 in adult vascular endothelium by tamoxifen injections. *Mettl7a1^ΔECKO^* were generated by breeding *Mettll7a1^fl/fl^* mice with *Cdh5-Cre/ERT2* mice with an inducible Cre recombinase under the control of the vascular endothelial cadherin (Cdh5) promoter. *Mettll7a1^fl/fl^ C57BL/6* mice were generated by inserting a loxP site at the promoter and downstream of exon 2 of Mettl7a1, respectively. **(c)** *Mettl7a1* mRNA expression was significantly reduced in carotid intima endothelial cells in tamoxifen-injected *Mettl7a1^ΔECKO^* compared to tamoxifen-injected *Cdh5-Cre/ERT2* mice (n = 6 for *Cdh5-Cre/ERT2* mice, n = 9 for *Mettl7a1^ΔECKO^* mice). \*\**p*≤0.01, *p* values were obtained using two-tailed Mann-Whitney U test using GraphPad Prism. Data were presented as mean±SD.

**Supplemental Figure 7.**
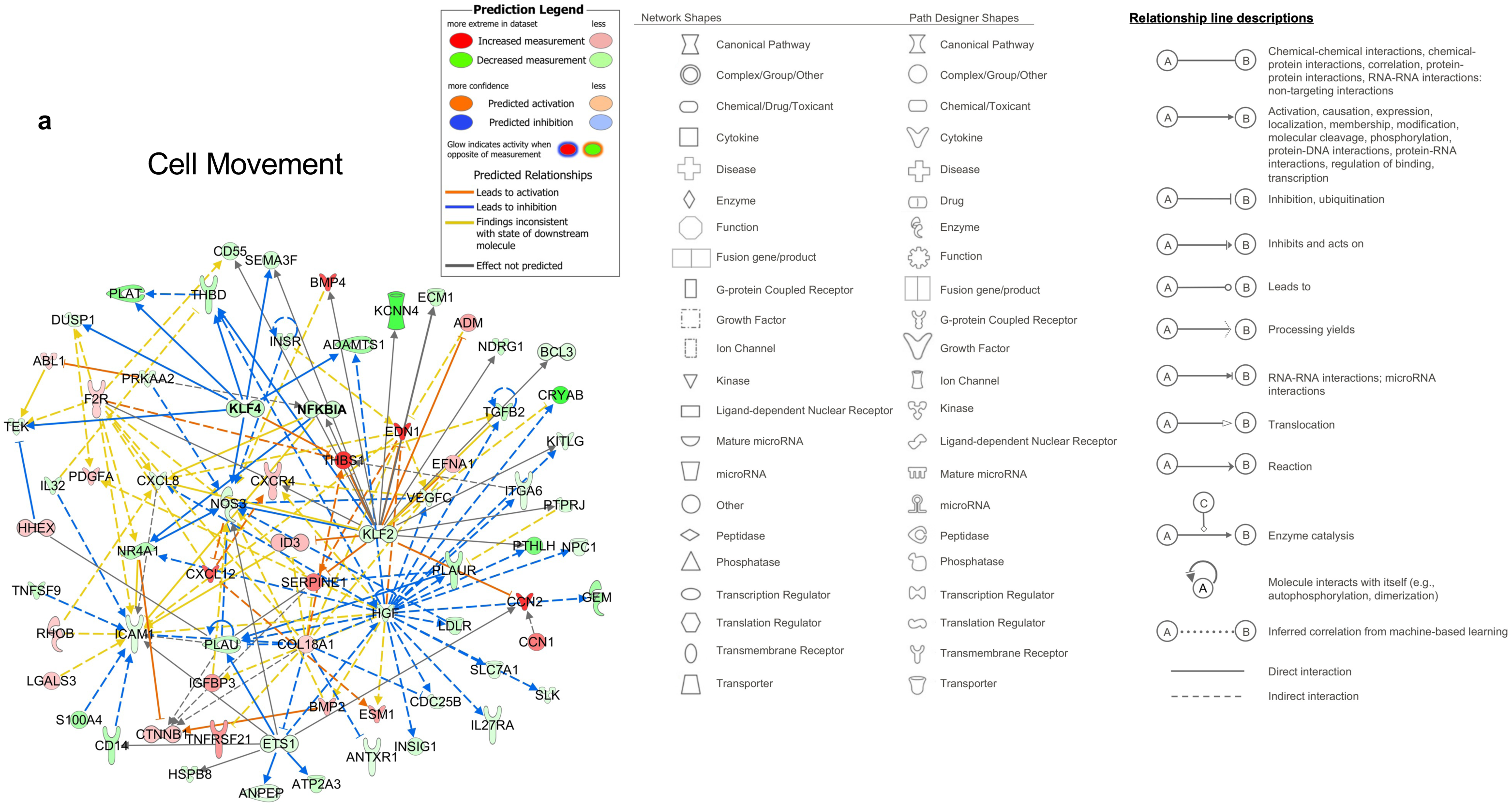
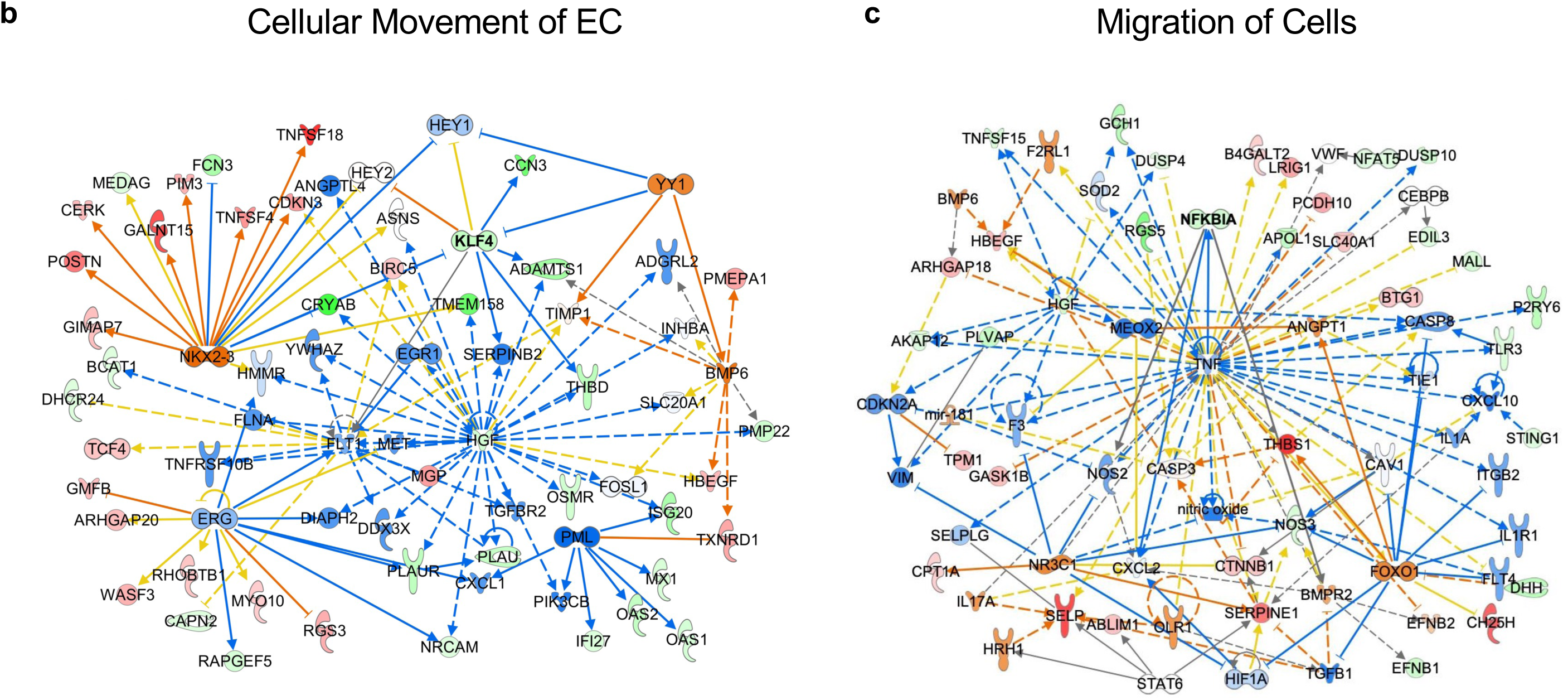
Ingenuity Pathway Analysis (IPA) of the METTL7A-mediated endothelial transcriptome identifies (a) Cell Movement, (b) Cell Movement of Endothelial Cells, and (c) Migration of Cells as top annotated biological functions in which KLF4 and/or NFKBIA are highly-connected hub genes. Endothelial genes downregulated by METTL7A knockdown are shown in green whereas genes upregulated by METTL7A knockdown are shown in red, as indicated by IPA legend which also describes the molecule types and regulatory relationships.

**Supplemental Figure 8.**
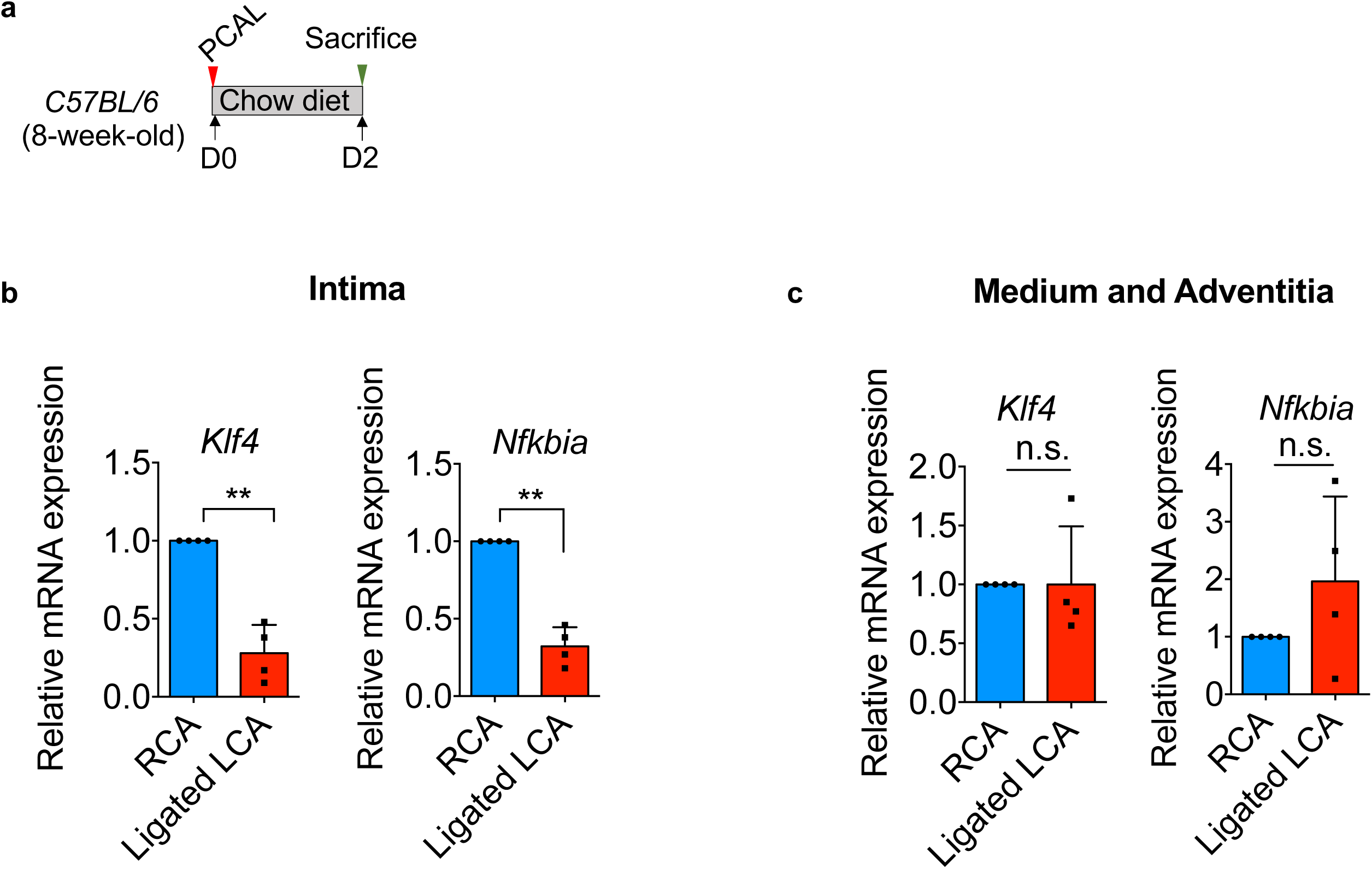
Acute disturbed flow *in vivo* significantly reduces endothelial expression of *Klf4* and *Nfkbia*. **(a)** Partial ligation was conducted in the left carotid artery of *C57BL/6* mice and carotid tissues were collected 48 hrs after the surgery. **(b)** *Klf4* and *Nfkbia* mRNAs were significantly reduced in the endothelium-enriched intima in the ligated left carotid artery compared to those in the right carotid artery **(c)** *Klf4* and *Nfkbia* mRNA expression in the carotid media and adventitia was not affected by the acute disturbed blood induced by partial carotid ligation. (n = 4 biological repeats, \*\**p*≤0.01 and n.s. indicates non-significance (*p*>0.05)). *p* values were obtained using two-tailed Student’s t-test using GraphPad Prism. Data were presented as mean±SD.

**Supplemental Figure 9.**
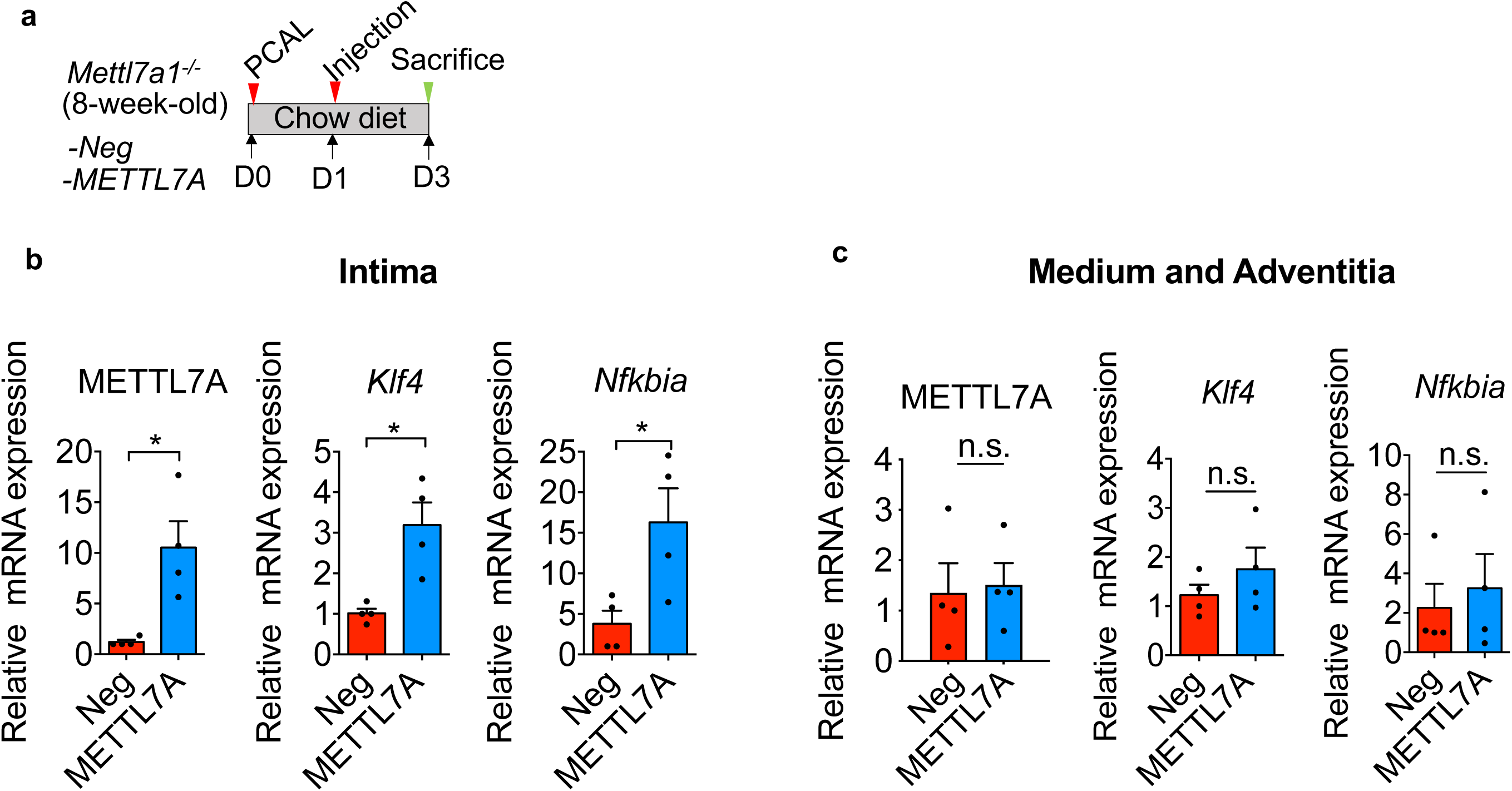
Endothelial METTL7A overexpression upregulates *Klf4* and *Nfkbia* intimal expression in *Mettl7a1^-/-^* mice. **(a)** Nanoparticles encapsulating CDH5-Ctrl (Neg) or CDH5-METTL7A (METTL7A) plasmids were intravenously administered to *Mettl7a1^-/-^* mice subjected to partial carotid ligation (PCAL). **(b)** Single intravenous injection of nanoparticles encapsulating CDH5-METTL7A plasmids (2 days after PCAL) significantly restored *Klf4* and *Nfkbia* mRNA expression in the endothelium-enriched intima in the ligated carotid artery of *Mettl7a1^-/-^* mice, compared to those receiving nanoparticles encapsulating CDH5-Ctrl plasmids. **(c)** Expression of METTL7A, *Klf4*, and *Nfkbia* in the media/adventitia in the ligated carotid artery of *Mettl7a1^-/-^* mice was not affected by the injection of nanoparticles encapsulating CDH5-METTL7A plasmids (n = 4 biological repeats). *p* values were obtained using two-tailed Student’s t-test using GraphPad Prism. \**p*≤0.05 and n.s. indicates non-significance (*p*>0.05). Data were presented as mean±SD.

**Supplementary Figure 10.**
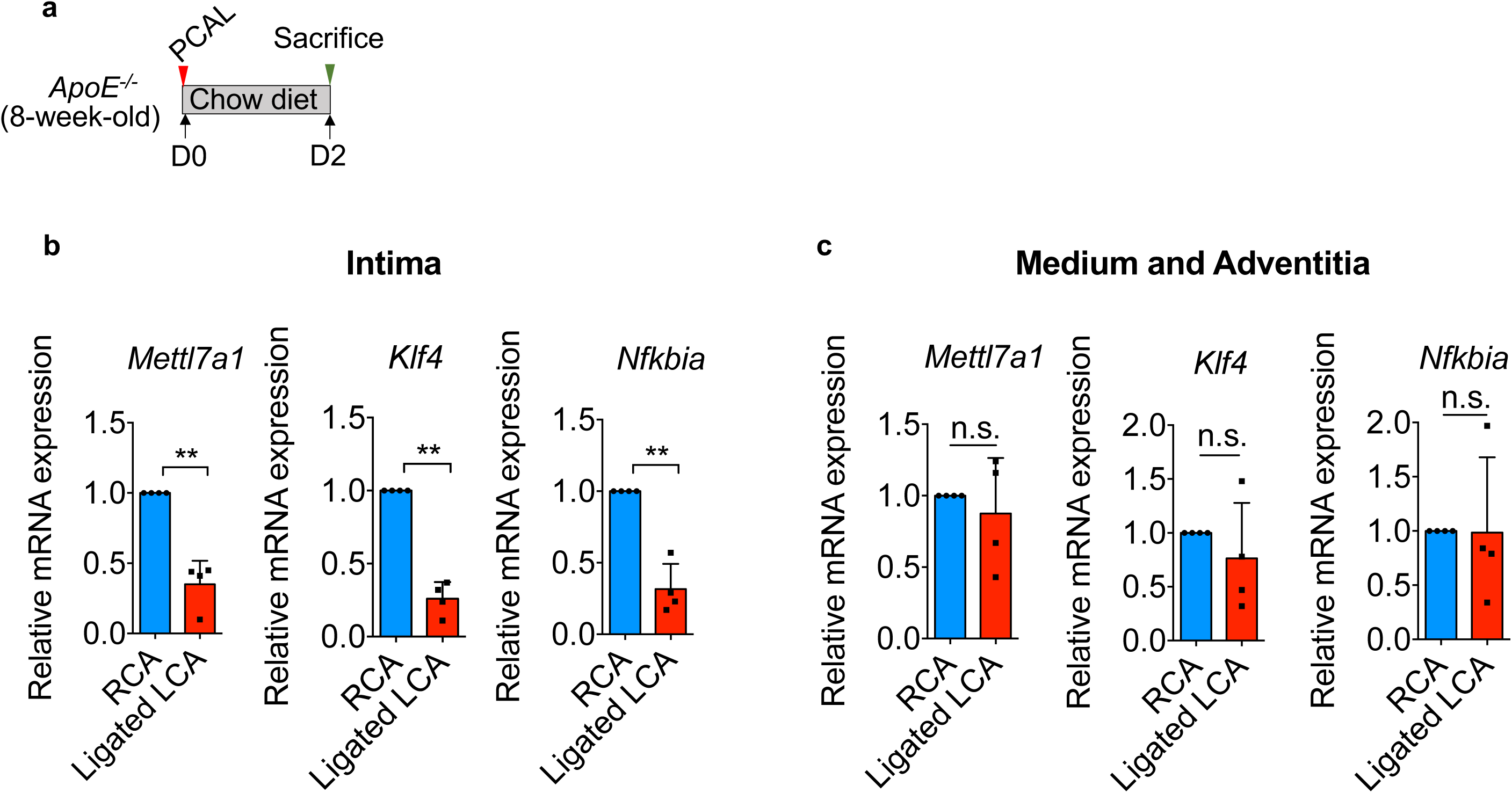
Endothelial expression of *Mettl7a1*, *Klf4*, and *Nfkbia* is significantly reduced by disturbed flow in the ligated left carotid artery in *ApoE^−/−^* mice. (a) Partial carotid ligation (PCAL) was performed to induce disturbed flow in the left carotid artery (LCA) of *ApoE^−/−^* mice. (b) 48-hr acute disturbed flow in the partially-ligated left carotid artery markedly reduced the transcripts of *Mettl7a1*, *Klf4*, and *Nfkbia* in endothelium-enriched intima. (c) 48-hr acute disturbed flow in the partially-ligated left carotid artery had no effect on the transcripts of *Mettl7a1*, *Klf4*, and *Nfkbia* in the media and adventitia (n = 4 biological repeats). *p* values were obtained using two-tailed Student’s t-test using GraphPad Prism. \*\**p*≤0.01 and n.s. indicates non-significance (*p*>0.05). Data were presented as mean±SD.

**Supplemental Table 1:**
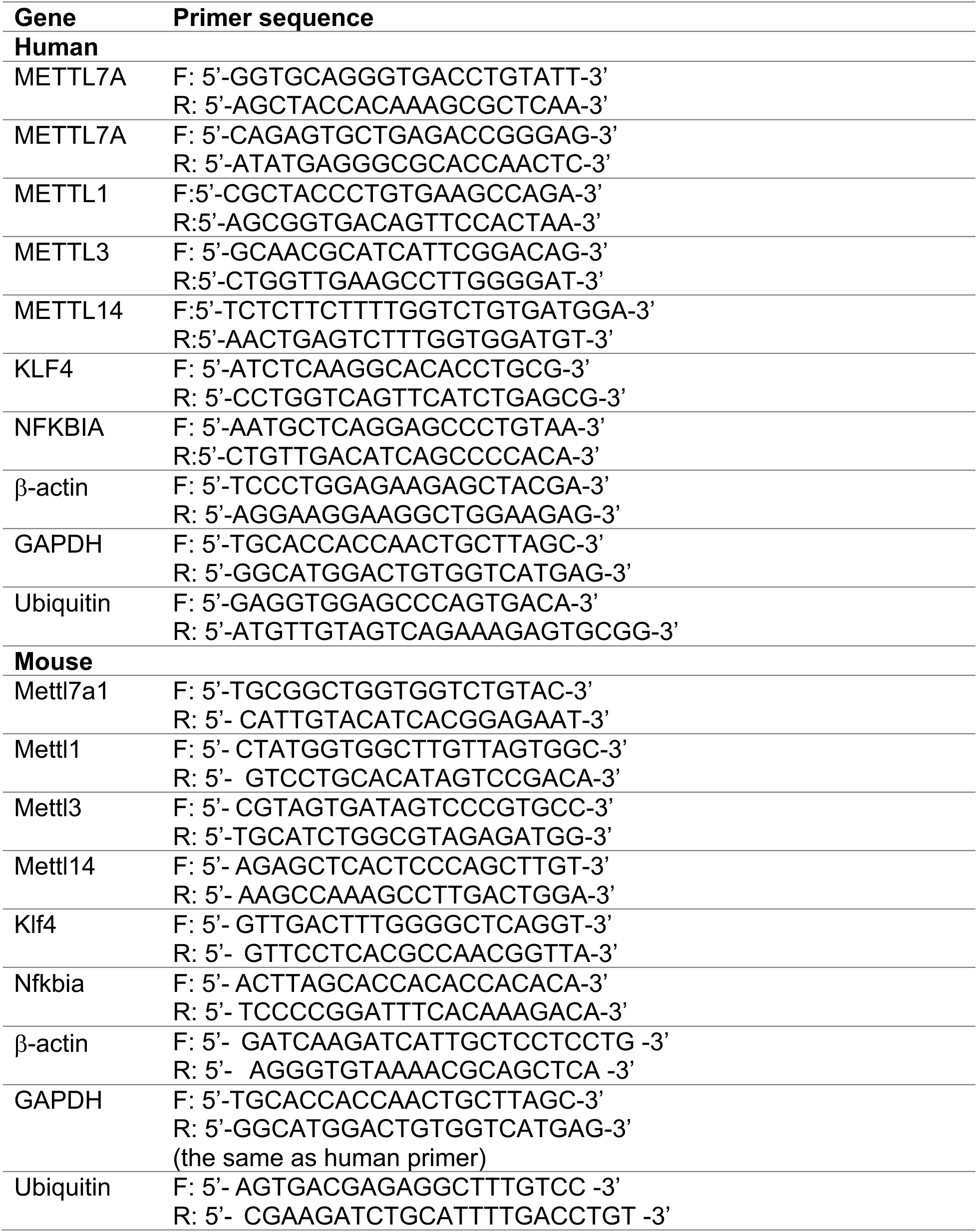
Primer List.

**Supplemental Table 2:** Top IPA-predicted upstream transcriptional regulators in the METTL7A-mediated endothelial transcriptome and their mRNA levels responding to METTL7A knockdown.

| Upstream regulator | Experimental Log Ratio<br>(UF+siMETTL7A/UF+siCtrl) |
| --- | --- |
| KLF4 | -1.149 |
| NFKBIA | -0.891 |
| MEF2C | -0.747 |
| KLF2 | -0.729 |
| ETS1 | -0.528 |
| FOXC1 | 0.365 |

**Supplemental Table 3:** A list of 115 endothelial genes, the mRNA of which are bound by METTL7A and regulated by METTL7A knockdown.

| Number | Gene | Experimental Log Ratio<br>(UF+siMETTL7A/UF+siCtrl) |
| --- | --- | --- |
| 1 | TPM1 | 0.481 |
| 2 | TPM4 | -1.450 |
| 3 | IL32 | -0.925 |
| 4 | MAN1A1 | -0.591 |
| 5 | PDGFB | 0.947 |
| 6 | RPS6KA2 | 0.452 |
| 7 | HMGN2 | 0.528 |
| 8 | SYNPO | -0.406 |
| 9 | HMGCS1 | -0.728 |
| 10 | KLF4 | -1.149 |
| 11 | PAX8-AS1 | -0.706 |
| 12 | LUCAT1 | -2.158 |
| 13 | TXNRD1 | 0.654 |
| 14 | CKB | -0.844 |
| 15 | SORT1 | 0.557 |
| 16 | CIRBP | 0.531 |
| 17 | DOCK5 | -0.771 |
| 18 | FZD8 | 0.985 |
| 19 | ESYT2 | -0.331 |
| 20 | HEG1 | -1.059 |
| 21 | EFEMP1 | 0.959 |
| 22 | SAT1 | 0.506 |
| 23 | MTUS1 | 0.575 |
| 24 | HPCAL1 | -0.489 |
| 25 | SCD | -0.539 |
| 26 | SCML1 | -0.484 |
| 27 | KLHL13 | 0.661 |
| 28 | PKM | -0.407 |
| 29 | RNU11 | -1.149 |
| 30 | MLLT1 | 0.454 |
| 31 | RNU5B-1 | -1.776 |
| 32 | PLAUR | -1.226 |
| 33 | HSPA1A | -1.139 |
| 34 | METTL7A | -1.139 |
| 35 | RNU1-27P | -5.658 |
| 36 | HSP90AA1 | -0.331 |
| 37 | PMP22 | -0.938 |
| **38** | **SELP** | **1.347** |
| **39** | **SEMA6B** | **0.332** |
| **40** | **SORBS2** | **0.636** |
| **41** | **VWF** | **0.531** |
| **42** | **ANPEP** | **-0.389** |
| **43** | **ADAMTS1** | **-1.779** |
| **44** | **PODXL** | **-0.892** |
| **45** | **MTHFD1L** | **0.449** |
| **46** | **ELOVL1** | **-0.338** |
| **47** | **KLF15** | **0.893** |
| **48** | **ID3** | **0.508** |
| **49** | **BCAR1** | **0.446** |
| **50** | **CPT1A** | **0.356** |
| **51** | **CREM** | **-0.599** |
| **52** | **SLC2A3** | **-0.894** |
| **53** | **RHOB** | **0.389** |
| **54** | **RHOC** | **-0.387** |
| **55** | **SLC12A2** | **-0.457** |
| **56** | **TRIB3** | **-0.420** |
| **57** | **CLEC14A** | **0.547** |
| **58** | **KCTD20** | **-0.350** |
| **59** | **ARHGAP27** | **-0.382** |
| **60** | **ATP1A1** | **-0.361** |
| **61** | **ATP1B1** | **-0.699** |
| **62** | **NFAT5** | **-0.501** |
| **63** | **IL4R** | **-0.367** |
| **64** | **B4GALT2** | **0.353** |
| **65** | **WSB1** | **0.599** |
| **66** | **PLK2** | **0.770** |
| **67** | **ATP6V0A1** | **-0.560** |
| **68** | **ADAM19** | **-1.562** |
| **69** | **INSIG1** | **-0.880** |
| **70** | **GAS6** | **0.762** |
| **71** | **CD55** | **-0.979** |
| **72** | **IRAK1** | **0.357** |
| **73** | **GBE1** | **-0.555** |
| **74** | **TNRC6B** | **-0.478** |
| **75** | **GCH1** | **-1.369** |
| **76** | **WNK1** | **-0.467** |
| **77** | **MTRNR2L12** | **-1.373** |
| **78** | **BGN** | **0.341** |
| **79** | **ITPR3** | **-0.758** |
| **80** | **SLC38A1** | **-0.529** |
| 81 | CMTM6 | -0.374 |
| 82 | MAGED4B | -2.151 |
| 83 | PTPRF | 0.370 |
| 84 | DHCR24 | -0.515 |
| 85 | FAM107B | 0.379 |
| 86 | LYPD5 | -2.768 |
| 87 | NFKBIA | -0.891 |
| 88 | MTATP8P1 | -0.518 |
| 89 | MTATP6P1 | -0.518 |
| 90 | HNRNPA0 | 0.309 |
| 91 | EEF1A1P9 | -0.557 |
| 92 | ATP13A3 | -0.903 |
| 93 | BLOC1S5-TXNDC5 | 0.844 |
| 94 | TCF4 | 0.443 |
| 95 | CALD1 | 0.464 |
| 96 | CH25H | 1.178 |
| 97 | TTYH3 | -0.452 |
| 98 | ODC1 | -0.474 |
| 99 | ZMIZ1 | -0.734 |
| 100 | SNHG19 | 0.636 |
| 101 | RUNX1T1 | 0.473 |
| 102 | RFX2 | 0.809 |
| 103 | LGALS9 | -0.664 |
| 104 | ABLIM1 | 0.461 |
| 105 | THBS1 | 1.539 |
| 106 | SH3PXD2B | -0.440 |
| 107 | EFNA1 | 0.497 |
| 108 | MTA1 | 0.407 |
| 109 | TKT | 0.331 |
| 110 | SERPINE1 | 1.015 |
| 111 | RNY1 | -1.057 |
| 112 | SNORA49 | 0.464 |
| 113 | LTBP2 | 0.488 |
| 114 | CSMD1 | 0.900 |
| 115 | TM4SF1 | 0.351 |

